# Non-linear sequence similarity between the *Xist* and *Rsx* long noncoding RNAs suggests shared functions of tandem repeat domains

**DOI:** 10.1101/630475

**Authors:** Daniel Sprague, Shafagh A. Waters, Jessime M. Kirk, Jeremy R. Wang, Paul B. Samollow, Paul D. Waters, J. Mauro Calabrese

## Abstract

The marsupial inactive X chromosome expresses a long noncoding RNA (lncRNA) called *Rsx* that has been proposed to be the functional analogue of eutherian *Xist*. Despite the possibility that *Xist* and *Rsx* encode related functions, the two lncRNAs harbor no linear sequence similarity. However, both lncRNAs harbor domains of tandemly repeated sequence. In *Xist*, these repeat domains are known to be critical for function. Using k-mer based comparison, we show that the repeat domains of *Xist* and *Rsx* unexpectedly partition into two major clusters that each harbor substantial levels of non-linear sequence similarity. *Xist* Repeats B, C and D were most similar to each other and to *Rsx* Repeat 1, whereas *Xist* Repeats A and E were most similar to each other and to *Rsx* Repeats 2, 3, and 4. Similarities at the level of k-mers corresponded to domain-specific enrichment of protein-binding motifs. Within individual domains, protein-binding motifs were often enriched to extreme levels. Our data support the hypothesis that *Xist* and *Rsx* encode similar functions through different spatial arrangements of functionally analogous protein-binding domains. We propose that the two clusters of repeat domains in *Xist* and *Rsx* function in part to cooperatively recruit PRC1 and PRC2 to chromatin. The physical manner in which these domains engage with protein cofactors may be just as critical to the function of the domains as the protein cofactors themselves. The general approaches we outline in this report should prove useful in the study of any set of RNAs.

## Introduction

The sex chromosomes of therian (eutherian and metatherian) mammals evolved from a pair of identical autosomes after the split of therian and monotreme mammals from their most recent common ancestor. Since that divergence, the Y chromosome has lost the large majority of its protein coding genes, creating a gene dosage imbalance between XY males and XX females. Part of the system that compensates this imbalance is a process known as X-chromosome Inactivation (XCI). Initiated early during female development, XCI results in the transcriptional silencing of one X chromosome in each somatic cell in female mammals. In eutherians, XCI is mediated by a long non-coding RNA (lncRNA) called *Xist* (Balaton et al., 2018; Brockdorff, 2018; da Rocha and Heard, 2017; Sahakyan et al., 2018).

The silencing function of *Xist* is thought to be mediated by the concerted action of several domains of tandemly repeated sequence that are interspersed throughout its length. These repeat domains harbor binding sites for distinct subsets of proteins that, through incompletely understood mechanisms, help *Xist* achieve different aspects of its function. “Repeat A” binds proteins called SPEN and RBM15, and is required for the stabilization of spliced *Xist* RNA, and for *Xist* to silence actively transcribed regions of the X chromosome (Chu et al., 2015; Engreitz et al., 2013; Hoki et al., 2009; McHugh et al., 2015; Moindrot et al., 2015; Monfort et al., 2015; Patil et al., 2016; Royce-Tolland et al., 2010; Wutz et al., 2002). “Repeat B”, and at least a portion of “Repeat C”, bind HNRNPK to recruit the Polycomb Repressive Complex 1 (PRC1) to the inactive X chromosome (Almeida et al., 2017; Pintacuda et al., 2017). “Repeat E” binds many proteins, including CIZ1, and is required for the stable association of *Xist* with X-linked chromatin and for the sustained recruitment of Polycomb Repressive Complex 2 (PRC2) to the inactive X (Ridings-Figueroa et al., 2017; Smola et al., 2016; Sunwoo et al., 2017).

Intriguingly, metatherians (marsupials) may have convergently evolved their own lncRNA, *Rsx*, to mediate XCI in XX females (Grant et al., 2012). *Rsx* shares no linear sequence similarity with *Xist* and is located in a different syntenic block on the marsupial X. Nevertheless, *Rsx* shares a number of surprising similarities with *Xist*. Both *Xist* and *Rsx* are expressed exclusively from the inactive X in females and are retained in the nucleus, forming what has been described as a “cloud-like” structure around their chromosome of synthesis. Moreover, both lncRNAs are spliced yet unusually long in their final processed form, and their expression correlates with the accumulation of histone modifications deposited by the Polycomb Repressive Complexes (PRCs) on the inactive X (Grant et al., 2012; Wang et al., 2014).

Studies performed over the last three decades indicate that *Xist* is required for normal XCI in eutherians (Balaton et al., 2018; Brockdorff, 2018; da Rocha and Heard, 2017; Sahakyan et al., 2018). Given the similarities between *Rsx* and *Xist*, it has been proposed that the marsupial *Rsx* is the functional analogue of *Xist* (Grant et al., 2012). While this hypothesis has yet to be directly tested, expression of an *Rsx* transgene on a mouse autosome does, to a certain extent, induce local gene silencing, supporting the notion that *Rsx* harbors *Xist*-like function (Grant et al., 2012).

Despite their lack of linear sequence similarity, *Xist* and *Rsx* both harbor long, internal domains of tandemly repeated sequence (Grant et al., 2012; Johnson et al., 2018). We recently discovered that evolutionarily unrelated lncRNAs that encode similar functions often harbor non-linear sequence similarity in the form of k-mer content, where a k-mer is defined as all possible combinations of a nucleotide substring of a given length k (Kirk et al., 2018). Below, we describe our use of k-mer based methods to investigate the possibility that the repeat domains in *Xist* and *Rsx* harbor non-linear sequence similarity that might be suggestive of shared function.

## Results

### Lack of linear sequence similarity between repeat domains in *Xist* and *Rsx*

*Xist* and *Rsx* are both notable for their domains of highly repetitive sequence, which can be identified by aligning each lncRNA to itself and visualizing the alignment data as a dot plot (File S1; (Rice et al., 2000)). In mouse *Xist*, the four major repetitive regions are referred to as Repeats A, B, C, and E (Figure 1A; (Brockdorff et al., 1992)). Repeats A, B, and E are conserved in eutherian mammals, whereas Repeat C appears to be specific to murid rodents (Figure 1C; (Nesterova et al., 2001; Yen et al., 2007)). In human *Xist*, the four major repetitive regions are referred to as Repeats A, B, D, and E (Figure 1B; (Brown et al., 1992)). Relative to mouse, human Repeat B is comprised of two shorter Repeat B-like regions that appear to have been disrupted by insertion (Figure 1B; (Nesterova et al., 2001; Yen et al., 2007)). Human Repeat D is comprised of eight core repeats flanked by several additional repeats that exhibit partial similarity to its core (Figure 1B; (Brown et al., 1992; Nesterova et al., 2001; Yen et al., 2007)). While Repeat D is absent in murid rodents (Figure 1C), Repeat D-like sequence appears in many other mammals (Figure S1; File S2; (Nesterova et al., 2001; Yen et al., 2007)).

**Figure 1.**
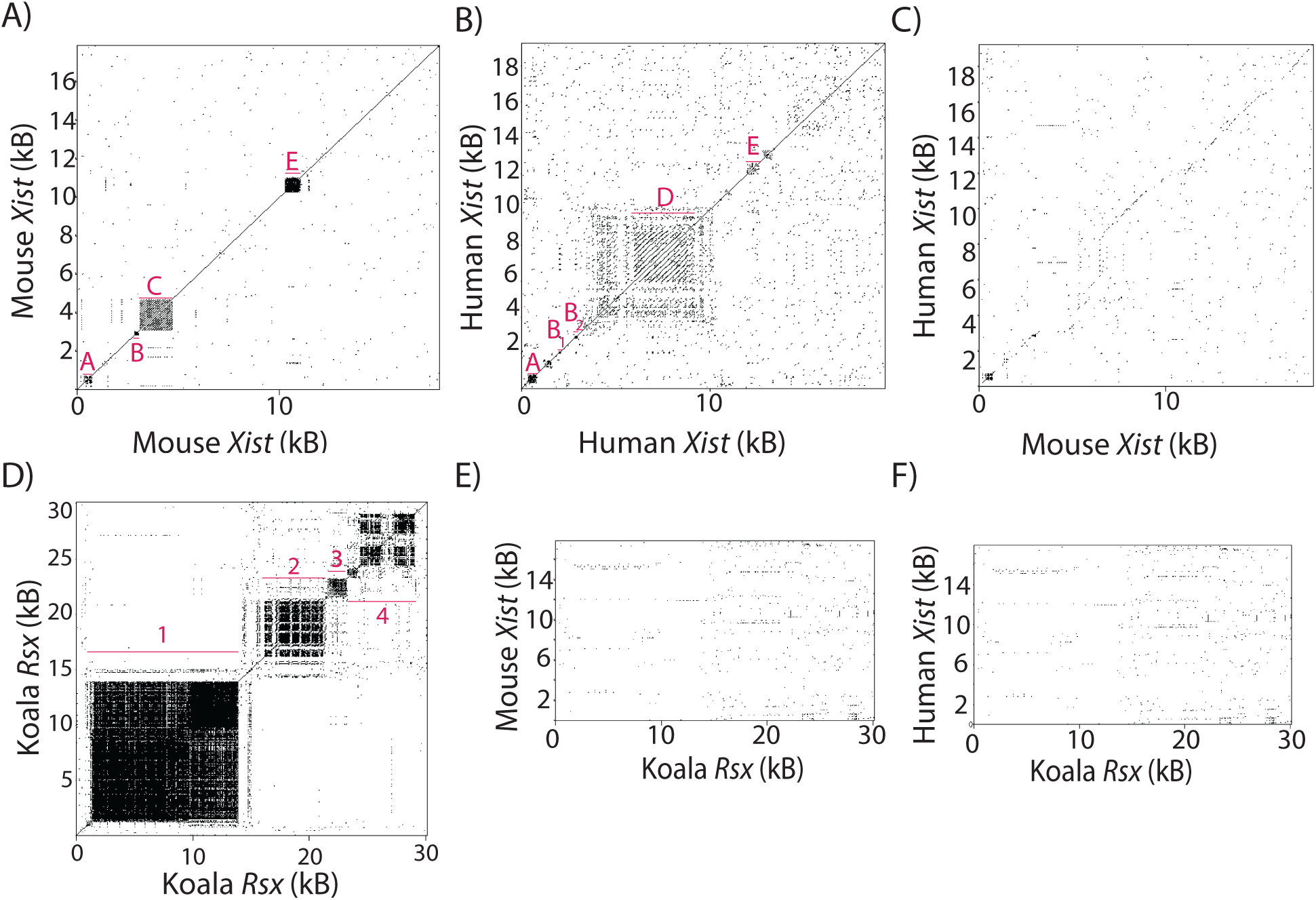
Lack of linear sequence similarity between repeat domains in *Xist* and *Rsx*. **(A-F)** Dot plots comparing mouse and human *Xist* and *Rsx* to themselves and to each other. The location of repeat domains in all three lncRNA-to-self plots are marked with red bars and names/numbers.

In contrast to *Xist*, which is mostly comprised of non-repetitive sequence, nearly all of the sequence in *Rsx* can be assigned to one of four repetitive domains (Figure 1D; (Johnson et al., 2018)). Here, we refer to the repetitive domains in *Rsx* as Repeats 1 through 4. It has been suggested that *Rsx* Repeat 1 is functionally analogous to *Xist* Repeat A, because both repeats are the first to occur in each lncRNA, and because both repeats contain GC-rich elements (Grant et al., 2012; Johnson et al., 2018). Beyond this observation, little is known about the repetitive regions in *Rsx* and how they might relate to those in *Xist*. Hypothesizing that the repeat domains in *Xist* and *Rsx* recruit similar subsets of proteins, we expected that dot plots comparing the sequence of *Xist* to the sequence of *Rsx* would reveal regions of sequence similarity. However, this was not the case (Figure 1E, F), nor was significant similarity between *Xist* and *Rsx* detected using BLASTN or the hidden-markov based nhmmer (Altschul et al., 1990; Wheeler and Eddy, 2013).

### Non-linear similarity between repeat domains in *Xist* and *Rsx*

We hypothesized that sequence similarity between *Xist* and *Rsx* might become apparent using an algorithm we recently developed to detect sequence similarity between evolutionarily unrelated lncRNAs (Kirk et al., 2018). In our algorithm, called SEEKR (<u>Se</u>quence <u>e</u>valuation through <u>k</u>-mer representation), groups of lncRNAs are compared to each other by counting the number of occurrences of each k-mer of a given length k in each lncRNA, then normalizing k-mer counts by the length of the lncRNA in question, and finally calculating a z-score for each k-mer in each lncRNA. The list of z-scores for each k-mer in a lncRNA is referred to as its “k-mer profile” and represents the abundance of each k-mer in the lncRNA relative to the abundance of each k-mer in the other lncRNAs that were analyzed as part of the group. In SEEKR, k-mer profiles from lncRNAs of interest are compared to each other using Pearson’s correlation. We previously demonstrated that SEEKR can be used to quantify the similarity between any number of lncRNAs regardless of their evolutionary relationships or differences in their lengths, and that similarities in k-mer profiles correlated with lncRNA protein binding potential, subcellular localization, and *Xist*-like repressive activity. A major strength of SEEKR is that it ignores positional information in similarity calculations, allowing it to quantify non-linear sequence relationships (Kirk et al., 2018).

In order to compare *Xist* and *Rsx* via SEEKR, we calculated the k-mer profile at k=4 of individual repeat domains in mouse *Xist* and koala *Rsx*, using all mouse lncRNAs from GENCODE as a background set to derive the mean and standard deviation of the counts for each k-mer (Derrien et al., 2012). The mechanisms through which *Xist* functions have been most extensively studied in mouse (Sahakyan et al., 2018). For this reason, we primarily used the repetitive regions from mouse *Xist* as search features in this work. However, because of the conservation of Repeat D-like domains in non-murid eutherian mammals (Figure S1; (Nesterova et al., 2001; Yen et al., 2007)), we also included the sequence of human *Xist* Repeat D in our analyses. We used the sequence from koala *Rsx* as our exemplar, owing to the high-quality of the koala genome build relative to builds from other marsupials (Johnson et al., 2018).

In our previous work, we found that SEEKR performed best when the length of the lncRNA or lncRNA fragment being studied was similar to 4^k, i.e. the total number of possible k-mers at k-mer length k. In tests of *Xist*-like repressive activity, we found that comparisons of lncRNAs using k-mer lengths of k≥7 underperformed relative to comparisons using smaller k-mer lengths, owing to the fact that most annotated lncRNAs are much less than 4^7 (16384) nucleotides long, and k-mer profiles of individual lncRNAs at k≥7 (≥16384 possible k-mers) are dominated by “0” values (Kirk et al., 2018). Based on this observation, and because Repeats A and B, two essential repetitive regions within *Xist* (Almeida et al., 2017; Hoki et al., 2009; Pintacuda et al., 2017; Royce-Tolland et al., 2010; Wutz et al., 2002), are each about 4^4 (256) nucleotides in length, we reasoned that k-mer profiles at k=4 (4^4=256 possible k-mers) would provide a reasonable estimate of sequence complexity for the repeats without being dominated by “0” values.

We also noted that relative to most lncRNAs, k-mer content in the repetitive regions of *Xist* and *Rsx* was skewed (Figure S2A). We therefore elected to log_2_-transform z-scores in k-mer profiles prior to comparison via Pearson’s correlation, recognizing that this transformation would reduce skew and allow us to evaluate similarity in the context of a log-linear scale (Figure S2B).

The individual repeat domains in *Xist* and *Rsx* vary substantially in terms of their length and sequence complexity. *Xist* repeats tend to be shorter and lower in overall complexity than repeats in *Rsx* (Figure 2A, B). Despite these differences, using SEEKR, we identified substantial levels of similarity between the repeat domains of *Xist* and *Rsx*. The Repeat A region of *Xist* was most similar to *Rsx* Repeat 4, exhibited a weak positive correlation with Repeat 2, and had negative correlations with *Rsx* Repeats 1 and 3 (Pearson’s r of 0.21, for Repeat A versus Repeat 4, respectively, and r of −0.02, 0.09, and −0.08 for Repeats 1, 2, and 3, respectively; Figure 2C). In contrast, *Xist* Repeat B was most similar to *Rsx* Repeat 1 and had negative correlations with *Rsx* Repeats 2 through 4 (Pearson’s r of 0.33 for Repeat B versus Repeat 1; Figure 2C). Repeat C, which is specific to murid rodents (i.e. it is not found in other eutherians), had no appreciable correlation with any *Rsx* repeat, whereas human Repeat D had positive correlations with *Rsx* Repeat 1 and 4 (r of 0.20, 0.12, respectively; Figure 2C). The k-mer profile of *Xist* Repeat E had positive correlations that increased progressively in *Rsx* Repeats 2, 3, and 4. (Pearson’s r of 0.15, 0.25, 0.40, respectively; Figure 2C).

**Figure 2.**
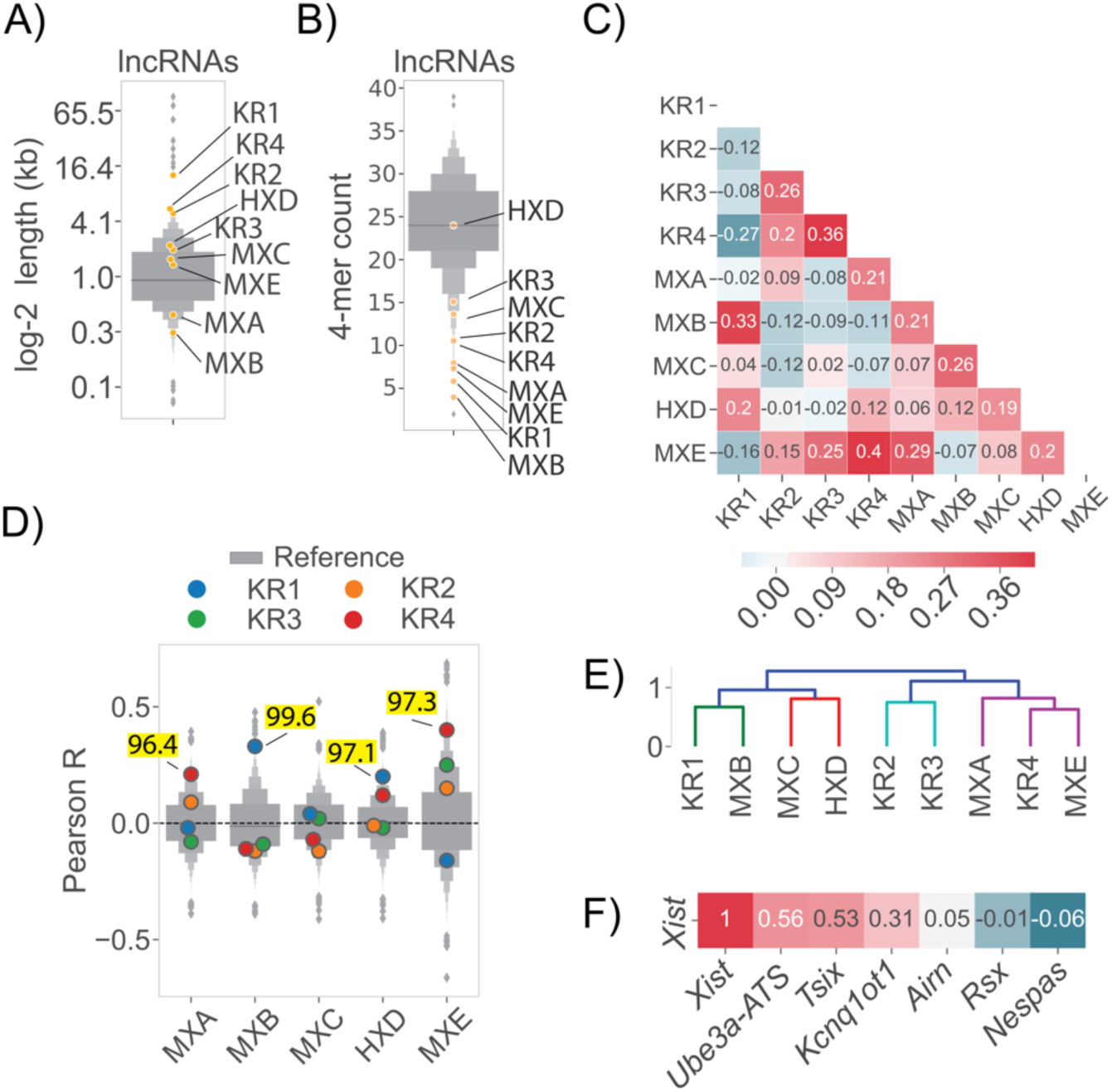
Non-linear similarity between repeat domains in *Xist* and *Rsx*. **(A)** Length of *Xist* and *Rsx* repeat domains and **(B)** sequence complexity estimated by the number of unique 4-mers that constitute 25% of the total 4-mer counts in a transcript, each relative to all other GENCODE M18 lncRNA transcripts. For panels A – E: “M”, “H”, and “K” signify mouse, human, and koala, respectively, “X” and “R” signify *Xist* and *Rsx*, respectively, and the final letter or number in each abbreviation signifies the repeat domain in question. **(C)** Correlation matrix displaying the Pearson’s r value derived from comparing k-mer profiles at k = 4 of each of the four repeat domains in koala *Rsx* (Repeats 1 – 4), the major repeats in mouse *Xist* (Repeats A, B, C, E), and human *Xist* Repeat D. The set of mouse lncRNAs from GENCODE was used to derive mean and standard deviation values for length normalized abundance of each k-mer. **(D)** Similarity of repeat domains in *Xist* and *Rsx* relative to all lncRNA transcripts in the mouse GENCODE M18 database (Derrien et al., 2012). Each subplot shows the distribution of Pearson’s r values describing the similarity between the *Xist* repeat in question and the set of GENCODE lncRNA transcripts. Similarities between *Xist* and *Rsx* that are above the 95^th^ percentile of similarity for all mouse lncRNAs are highlighted in yellow. **(E)** The correlation matrix in (A) subject to hierarchical clustering. Colors represent clusters for all descendent links beneath the first node in the dendrogram with distance less than 70% of the largest distance between all clusters. **(F)** SEEKR-derived similarity (in the form of Pearson’s r; (Kirk et al., 2018)) between full-length *Xist*, other *cis*-repressive lncRNAs in mouse, and koala *Rsx* (Johnson et al., 2018).

We sought to quantify the strength of the similarity between repeat domains in *Xist* and *Rsx* relative to other mouse lncRNAs. To do this, we used Pearson’s correlation to compare the k-mer profile of each *Xist* repeat domain to the k-mer profiles of the set of spliced GENCODE M18 mouse lncRNAs (Derrien et al., 2012). We compared this distribution of Pearson’s r values to the r value obtained when comparing each *Xist* repeat to each *Rsx* repeat.

This analysis revealed striking similarities between the repeat domains of *Xist* and *Rsx*. *Xist* Repeat B was more similar to *Rsx* Repeat 1 than it was similar to 99.6% of all lncRNAs (similarity ranked 65^th^ out of 17523 comparisons), despite the fact that the two repeats differ in length by ~50-fold (Figures 2A, D; Tables S1, S2). *Xist* Repeat A was more similar to *Rsx* Repeat 4 than it was similar to 96.4% of all other lncRNAs (its similarity ranked 626^nd^ out of 17523 comparisons), *Xist* Repeat D was more similar to *Rsx* Repeat 1 than it was similar to 97.1% of all other lncRNAs (its similarity ranked 515^th^ out of 17523 comparisons), and *Xist* Repeat E was more similar to *Rsx* Repeat 4 than it was similar to 97.3% of all other lncRNAs (its similarity ranked 467^th^ out of 17523 comparisons; Figure 2D; Table S1, S2). No other repeat domains in *Xist* and *Rsx* fell above the 95^th^ percentile in terms of their similarity to each other. Similar trends were observed when we used k-mer lengths k = 4, 5 and 6 for this analysis (Figure S3A).

Current models suggest that the tandem repeats in *Xist* have distinct functions (Balaton et al., 2018; Brockdorff, 2018; da Rocha and Heard, 2017; Sahakyan et al., 2018). Thus, we were surprised to find that the repeat domains within *Xist* also exhibited high levels of similarity to each other (Table S1, S2). Repeat A was more similar to Repeat E than it was similar to 99.6% of all lncRNAs. Likewise, Repeats B and C were more similar to each other than they were similar to 97.8% and 99.6% of all other lncRNAs, respectively. Finally, Repeats C and D were more similar to each other than they were similar to 97.0% and 96.2% of all other lncRNAs, respectively (Table S1, S2).

The similarities between specific domains of *Xist* and *Rsx* were also evident in an unsupervised hierarchical cluster of the matrix from Figure 2C. *Xist* Repeat B and *Rsx* Repeat 1 formed a basal cluster which joined with a second basal cluster comprising *Xist* Repeat C and *Xist* Repeat D. *Rsx* Repeat 4 and *Xist* Repeat E formed a basal cluster that that joined with *Xist* Repeat A. This multilevel cluster (*Rsx* Repeat 4, *Xist* Repeat E, and *Xist* Repeat A) joined with another basal cluster comprising *Rsx* Repeats 2 and 3 (Figure 2E).

At k-mer length k = 4, Pearson’s correlation with and without log-transformation of k-mer z-scores, as well as Spearman’s correlation of non-transformed z-scores, detected similar relationships between *Xist* and *Rsx* repeat domains (Figure S4). While the similarities between individual domains were still evident at higher k-mer lengths, particularly when using Pearson’s correlation of log-transformed k-mer counts, the clustering patterns that we observed at k-mer length k = 4 began to dissolve (Figure S4). At high k-mer lengths, Spearman’s correlation was the least informative method of comparison, owing the large number of “zero” values that populate k-mer profiles at these lengths (Figure S4). Thus, to a certain extent, the similarities in the repeat domains of *Xist* and *Rsx* are detectable regardless of prior assumptions about log-linear, linear, and monotonic relationships between k-mer profiles. However, the most robust similarities are detected using Pearson’s correlation of log-transformed k-mer counts (Figure S4).

We observed that the similarities between *Xist* and *Rsx* were obscured when the k-mer profiles of the full-length lncRNAs were compared to each other (Pearson’s r of −0.01 for the comparison of full-length *Xist* to full-length *Rsx*; Figure 2F). This loss of similarity highlights the utility of domain-based similarity searches, particularly for lncRNAs whose functional domains may comprise a fraction of their overall length. The dissimilarity between k-mer profiles of full-length *Xist* and full-length *Rsx* likely stems from the fact that virtually all of *Rsx* is comprised of repetitive sequence domains that harbor limited k-mer diversity relative to the non-repetitive sequence of *Xist* (Figure 2B and compare Figures 1A-C to 1D).

### Sequence properties of *Xist* and *Rsx* repeat domains

Qualitative similarities between *Xist* and *Rsx* repeat domains were also revealed using MEME to visualize motifs that were enriched within individual domains (Bailey et al., 2009). In *Xist* Repeat A, MEME identified one motif comprised of short runs of G and C nucleotides and one motif most notable for runs of T nucleotides (Figure 3A). Similar patterns were seen in the motifs enriched in *Rsx* Repeat 4 (Figure 3B). The single motif from Repeat B was almost exclusively comprised of two tandemly arranged ‘GCCCC’ motifs, and motifs containing runs of ‘G’ and ‘C’ nucleotides could be seen in *Rsx* Repeat 1 (Figure 3A, B). The pyrimidine-rich runs that were characteristic of *Xist* Repeat E were also observed *Rsx* Repeat 4 (Figure 3A, B). *Rsx* Repeat 2 was unique in its enrichment of AAAG and GAAA motifs (Figure 3B).

**Figure 3.**
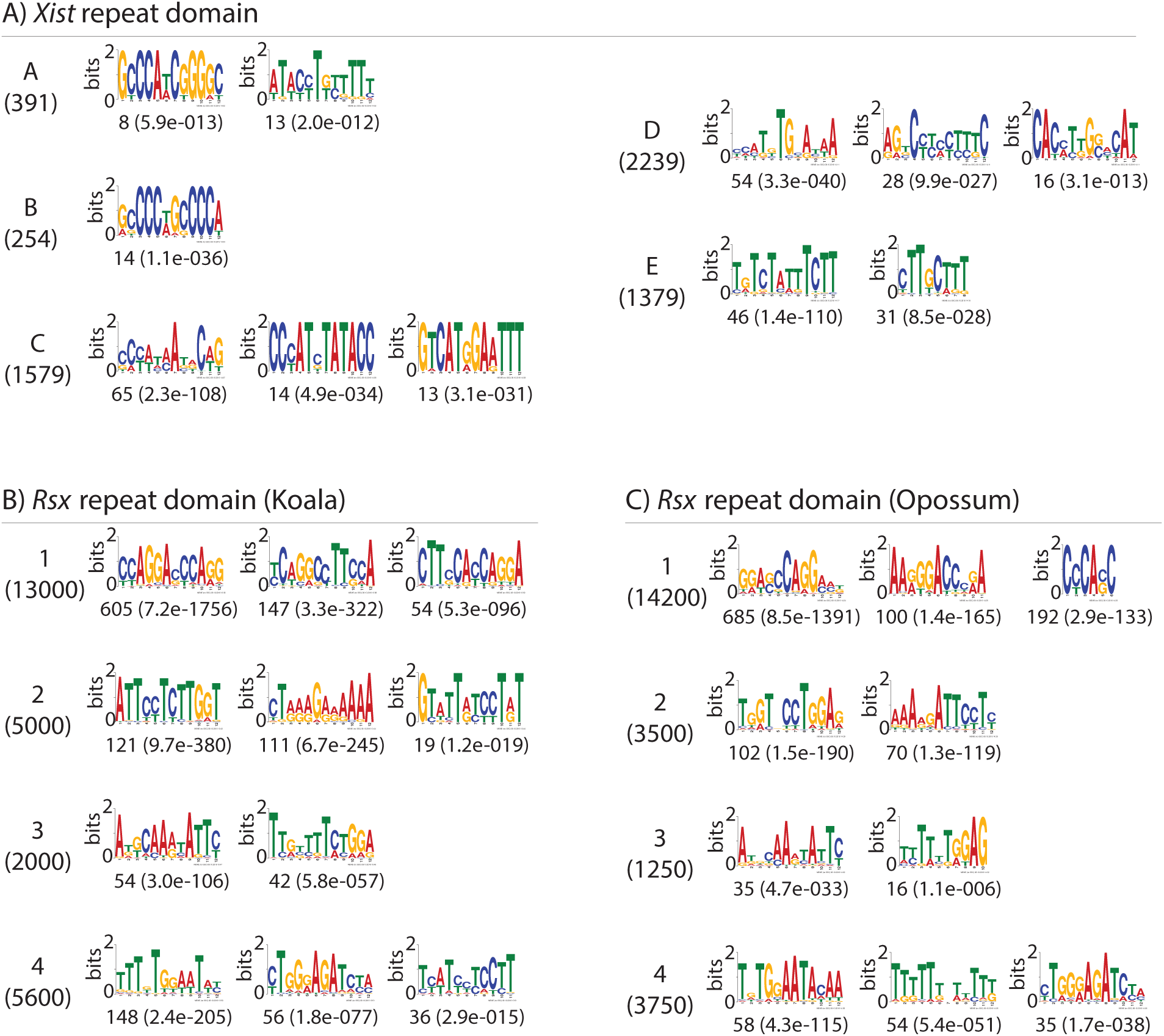
Motifs enriched in *Xist* and *Rsx* repeat domains. **(A-C)** The top three *de novo* motifs identified by MEME in *Xist* repeats (panel A; all repeats from mouse *Xist* except for Repeat D, which is the human sequence), and in the four repeats in koala (B) and opossum (C) *Rsx*. The length in nucleotides of each repeat is shown in parentheses below the repeat name. The number of matches to each motif, as well as the expectation-value for that number, is shown below each motif logo. Some repeats had less than three motifs detected by MEME.

Several of the repeat domains in *Xist* and *Rsx* could be distinguished by the presence of k-mers comprised of runs of individual nucleotides that extended for two or more consecutive positions (such as AA, CC, GG, or TT; File S1). Similar to enriched motifs, k-mers containing mononucleotide runs may function to recruit different subsets of RNA binding proteins (Dominguez et al., 2018; Ray et al., 2013). We therefore sought to quantify the enrichment of k-mers containing mononucleotide runs in the repeat domains of *Xist* and *Rsx*, reasoning that this analysis might provide insight into function.

Similar to what we observed in our motif analysis (Figure 3), *Rsx* Repeat 2 had the highest length-normalized abundance of polyA k-mers, followed closely by *Rsx* Repeat 3 (Figure 4A). Repeat B, which is only ~250 nucleotides long and is almost entirely comprised of polyC sequence, had the highest length-normalized abundance of polyC k-mers, followed by *Rsx* Repeat 1, and *Xist* Repeats C, D, and A (Figure 4B). Mouse Repeat A had the highest length-normalized abundance of polyG k-mers, followed by *Rsx* Repeats 1, 2, and 4 (Figure 4C). *Xist* Repeats A and E, as well as *Rsx* Repeats 3 and 4 had the highest length-normalized abundance of polyT k-mers, reflecting the high degree of SEEKR-detected similarity between these regions (Figure 4D). Similar trends were detected when we used k-mer lengths k=4, 5, and 6 for this analysis (Figure S3B). Thus, certain *Xist* and *Rsx* repeat domains share similarity in their overall k-mer profiles (Figure 2), in their enriched motifs (Figure 3), and in their enrichment in subsets of low-complexity k-mers that are comprised of mononucleotide runs (Figure 4). The repeat domains also harbor differences in sequence composition that are consistent with their lack of alignment via methods designed to detect linear sequence similarity (Figure 1).

**Figure 4.**
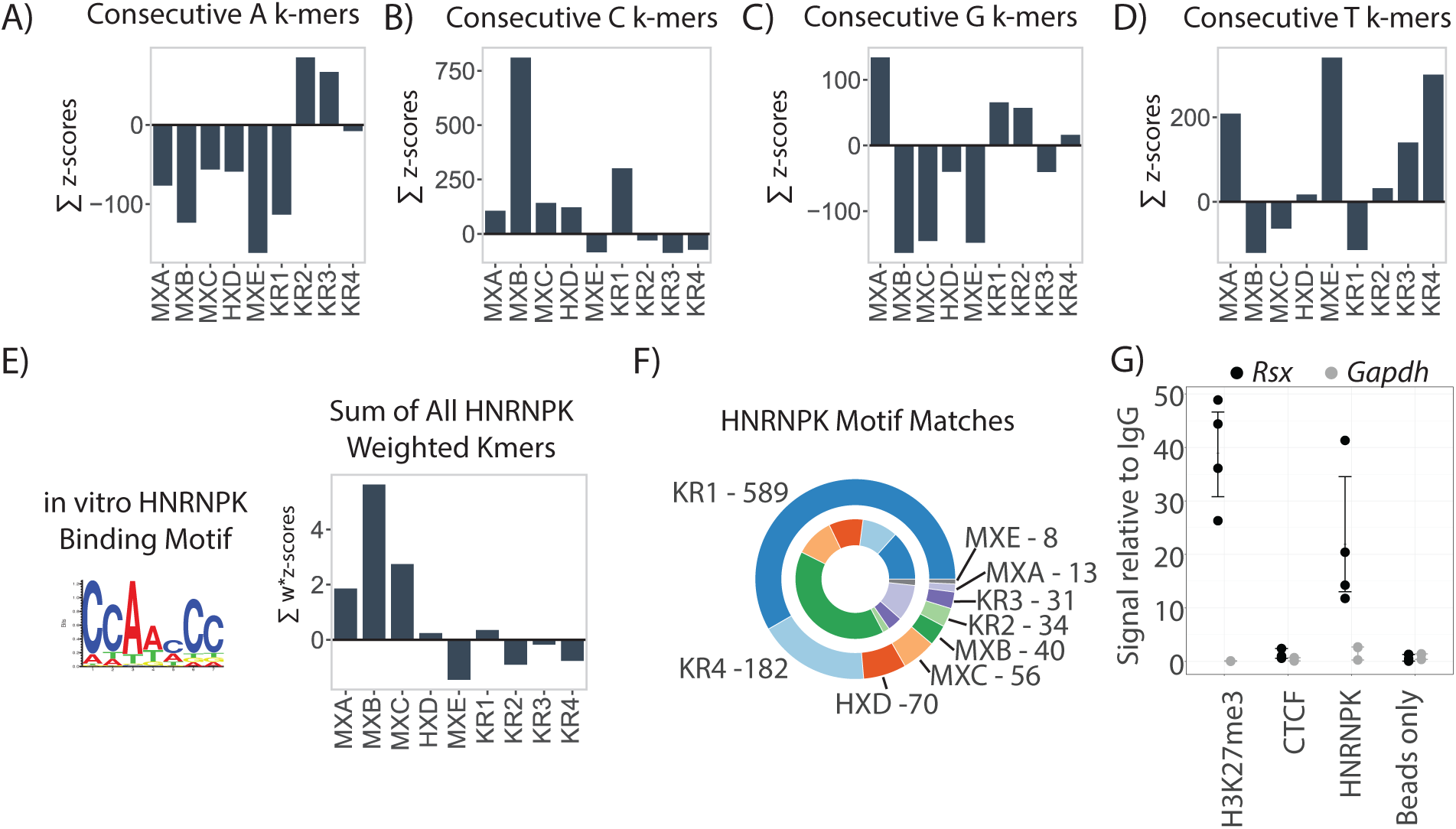
Mononucleotide runs and HNRNPK-binding motifs enriched in specific *Xist* and *Rsx* repeats. **(A-D)** The sum of z-scores in each repeat for k-mers containing consecutive (A) A, (B) C, (C) G, and (D) T nucleotides. For this analysis we defined “consecutive” as at least two consecutive nucleotides of the specified identity, and used k-mer length k = 5 (Methods). Repeat abbreviations as in Figure 2. **(E)** The sum of z-scores for all k-mers in each *Xist* and *Rsx* repeat domain after weighting the k-mers by the likelihood with which they fit the consensus HNRNPK-binding motif. Motif logo that describes the consensus HNRNPK-binding motif obtained from (Ray et al., 2013) is also shown. **(F)** Component arcs of outer circle indicate the proportion and number of HNRNPK-binding motif matches detected by FIMO (p<0.01) in each *Xist* and *Rsx* repeat domain. Component arcs of inner circle indicate the proportion of motif matches in each repeat domain normalized for domain length. Repeat abbreviations as in Figure 2. **(G)** *Rsx* enrichment relative to IgG control after RNA IP-qPCR in cultured fibroblasts from female *M. domestica*. For each antibody, left (black) is enrichment of *Rsx*, right (grey) is enrichment of *Gapdh*. The histone modification H3K27me3 is enriched on the inactive X in marsupials, so an association with *Rsx* was expected. CTCF has nanomolar affinity for RNA and along with “bead only”/no-antibody IP serves as a negative control demonstrating IP specificity (Kung et al., 2015). Dots represent values from replicate RNA IP experiments; error bars represent bootstrap 95% CI.

### HNRNPK-binding motifs are enriched in specific *Xist* and *Rsx* repeats

*Xist* Repeat B is known to bind a protein called HNRNPK, and this binding activity is essential for *Xist* to recruit PRC1 to the inactive X chromosome (Pintacuda et al., 2017). Given the quantitative and qualitative sequence similarities between *Xist* Repeat B and *Rsx* Repeat 1 (Figures 2 – 4), we sought to compare HNRNPK-binding potential between the two repeats using two conceptually distinct approaches. First, we weighted z-scores of individual k-mers in all *Xist* and *Rsx* repeat domains by the probability that the k-mer would occur in the Position-Weight-Matrix (PWM) describing the HNRNPK-binding motif from ((Ray et al., 2013); see PWM in Figure 4E). We then summed HNRNPK-scaled z-scores over each repeat, and plotted the results in a manner similar to Figure 4A-D. In this analysis, a positive sum indicates that k-mers matching the HNRNPK PWM occur more frequently in the domain in question than they occur in other lncRNAs in the GENCODE database.

On a length-normalized basis, *Xist* Repeats B, C, A, and D, in descending order, had positive sums of HNRNPK-scaled z-scores. Repeat 1 was the only repeat in *Rsx* to have a positive sum, perhaps consistent with a role in recruiting HNRNPK to *Rsx* (Figure 4E). The sum of HNRNPK-scaled z-scores in *Rsx* Repeat 1 was lower than the sums in *Xist* Repeats B, C, and A (Figure 4E), which might be taken as evidence that on a length-normalized basis, *Xist* Repeats B, C, and A have a higher density of k-mers that are likely to bind HNRNPK than *Rsx* Repeat 1 or any other *Rsx* repeat. However, at 13 kb in length, *Rsx* Repeat 1 is ~50 times longer than *Xist* Repeat B, and is over half of the length of full-length *Xist* itself (Brockdorff et al., 1992; Brown et al., 1992; Johnson et al., 2018). Thus, we also counted the absolute number of matches to HNRNPK-binding motifs in *Xist* and *Rsx* repeats. *Rsx* Repeat 1 had 15 times more matches to HNRNPK-binding motifs than did *Xist* Repeat B (589 matches in Repeat 1 compared to 40 matches in Repeat B; Figure 4F; (Bailey et al., 2009)). *Rsx* Repeat 4 also had a large number of matches to HNRNPK-binding motifs (182 matches), and human Repeat D and mouse Repeat C each had more HNRNPK-binding sites than Repeat B (70 and 56 matches, respectively, compared to 40 in Repeat B; Figure 4F). CLIP performed in mouse and human cells supports a direct association between HNRNPK and Repeat C and Repeat D, respectively (Figure S5; (Cirillo et al., 2016; Van Nostrand et al., 2016)). Collectively, these data support the ideas that mouse Repeat C and human Repeat D cooperate with Repeat B in recruiting HNRNPK to *Xist*, and suggest that *Rsx* Repeat 1, and to a lesser extent, *Rsx* Repeat 4, could also recruit HNRNPK to *Rsx*.

We next used RNA immunoprecipitation (RNA IP) followed by RT-qPCR to determine whether we could detect evidence of HNRNPK association with *Rsx*. In fibroblast cells derived from a female gray short-tailed opossum, *Monodelphis domestica*, we found that HNRNPK IP enriched for *Rsx* 20-fold over IgG control IPs (Figure 4G). This enrichment was similar to that seen for an IP using an antibody that detects histone H3-lysine27-trimethylation (H3K27me3), a modification known to be enriched on the opossum inactive X (Figure 4G; (Wang et al., 2014)). *Gapdh* mRNA was not enriched by IP of HNRNPK or H3K27me3 (Figure 4G). IP of CTCF, a protein that binds RNA with nanomolar affinity in a sequence non-specific manner, showed neither *Rsx* nor *Gapdh* enrichment (Figure 4G; (Kung et al., 2015)). Leaving out HNRNPK antibody prior to performing IP and qPCR also led to a loss of *Rsx* signal (“beads only” in Figure 4G). DNase-treated input RNA (no reverse transcription control) did not yield signal in qPCR assays, indicating DNase digestion prior to cDNA synthesis and qPCR proceeded to completion (not shown). These data support our computational analyses and suggest that HNRNPK associates with *Rsx* in marsupial cells.

### Conservation of repeat domains between koala and opossum *Rsx*

Considering that not all of the repeat domains in *Xist* exhibit conservation across eutherian mammals, we sought to determine whether or not the repeat domains in koala *Rsx* were conserved in another marsupial. *Rsx* was originally identified in opossum (Grant et al., 2012), but the most current assembly of the opossum genome (mondom5; (Casper et al., 2018)) harbors significant gaps within the sequence of *Rsx*.

To assemble a complete sequence of opossum *Rsx* for comparison to koala, we used Oxford Nanopore technology to sequence two Bacterial Artificial Chromosomes (BACs) that encompassed the opossum *Rsx* locus (VMRC18-839J22 and VMRC18-303M7). *De novo* assembly and polishing of sequence reads identified a single 235,139 base contig aligning to chrX that had on average a 0.5% error rate with the mondom5 assembly (File S3). Our assembly filled in 16620 bases of unannotated sequence in the *Rsx* locus, 361 bases of which were a part of the spliced *Rsx* lncRNA annotation from ((Grant et al., 2012); Table S3).

Alignment of this ungapped assembly of spliced opossum *Rsx* to koala *Rsx* revealed high levels of similarity between their repeat domains in a dot plot analysis (Figure 5A, B). This similarity could also be seen at the level of k-mers (Figure 5C, D), and by extraction of enriched motifs using MEME (Figure 3B, C). Opossum and koala, which are members of distantly related American and Australian marsupial families, respectively, diverged approximately 82 million years ago (Kumar et al., 2017). By comparison, mouse and human are separated by approximately 90 million years of evolution (Kumar et al., 2017). Repeat domains 1 through 4 in opossum and koala *Rsx* exhibited levels of sequence similarity that approximated or exceeded the similarity found between the repeat domains in mouse and human *Xist* (with the exception of Repeat C/Repeat D; Figure 5C-F). Thus, the repeat domains in *Rsx* appear to be at least as conserved between distantly related marsupials as the repeat domains in *Xist* are conserved among eutherians.

**Figure 5.**
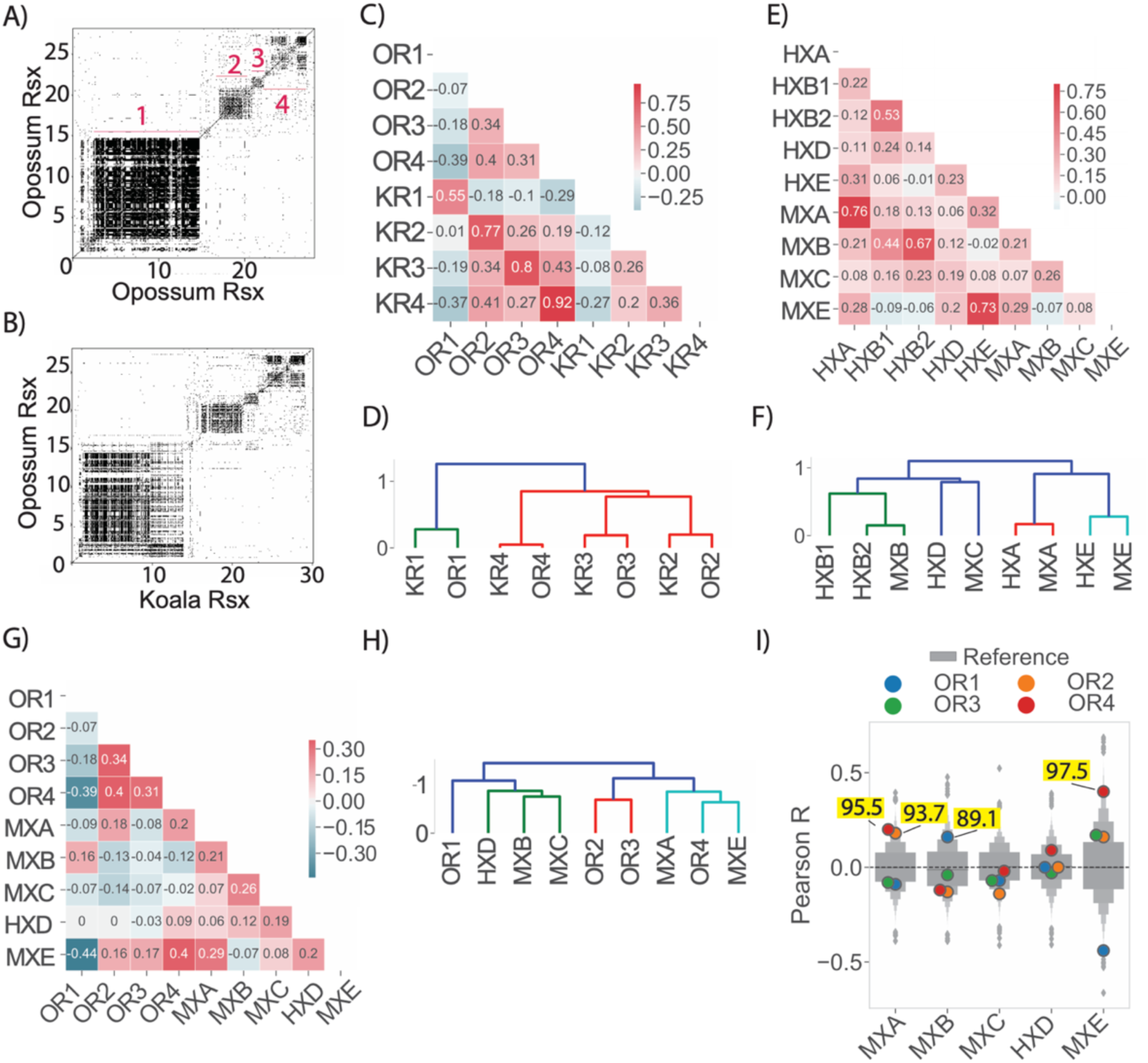
*Rsx* repeat domains are conserved between koala and opossum. **(A, B)** Dot plots of opossum *Rsx* aligned to (A) itself or (B) koala *Rsx*. **(C)** Similarity between repeat domains in koala and opossum *Rsx* as calculated in Figure 2C. **(D)** Hierarchical cluster of similarity values from (C). **(E)** Similarity between repeat domains in mouse and human *Xist* as calculated in Figure 2C. **(F)** Hierarchical clustering of similarity values from (E). **G)** Similarity between repeat domains in opossum *Rsx* and *Xist* repeat domains as calculated in Figure 2C. **(H)** Hierarchical cluster of similarity values from (G). **(I)** Percentiles for Pearson’s R for opossum *Rsx* repeat domains compared to each *Xist* repeat domain as in Figure 2D. Numbers mentioned in the body of the manuscript are highlighted in yellow.

Next, we compared the k-mer contents of repeat domains in *Xist* to the k-mer contents of repeat domains in opossum *Rsx*. We identified a level of similarity (Figure 5G, H, I) that mirrored the similarity we found between repeat domains in *Xist* and koala *Rsx* (Figure 2B, D, E). *Xist* Repeat A was most similar to opossum Repeats 2 and 4 (93.7^th^ and 95.5^th^ percentile relative to all other mouse lncRNAs, respectively); *Xist* Repeat B was most similar to opossum Repeat 1 (89.1^st^ percentile relative to all other lncRNAs); and *Xist* Repeat E was most similar to opossum Repeat 4 (97.5^th^ percentile relative to all other lncRNAs; Figure 5I). Thus, the major repeat domains in *Rsx* are conserved between opossum and koala, and the repeat domains in *Rsx* from both marsupials harbor k-mer contents similar those in repeat domains from mouse and human *Xist*.

### Multiple protein-binding motifs are enriched to extreme levels in *Xist* and *Rsx* repeat domains

We examined the extent to which *Xist* and *Rsx* repeat domains were enriched for sequence motifs known to recruit RNA binding proteins, hypothesizing that the patterns of enrichment might provide additional insight into similarities between the two lncRNAs. For this analysis, we downloaded PWMs for all mammalian RNA binding proteins available in the CISBP-RNA database (Ray et al., 2013), and for each PWM in each repeat, we quantified enrichment by weighting k-mer z-scores by the probability that the k-mer matched the PWM, then calculating the sum of those weights, as we did for the HNRNPK PWM in Figure 4E. To gauge the extent of enrichment relative to other mouse lncRNAs, we determined the percentile rank of the sum for each PWM in each repeat relative to the sums generated from the same PWM-weighting procedure performed on all mouse lncRNAs. We then hierarchically clustered repeat domains from *Xist* and *Rsx* based on the percentile ranks of motif enrichment for each domain. The results of these analyses are shown in Figure 6A.

**Figure 6.**
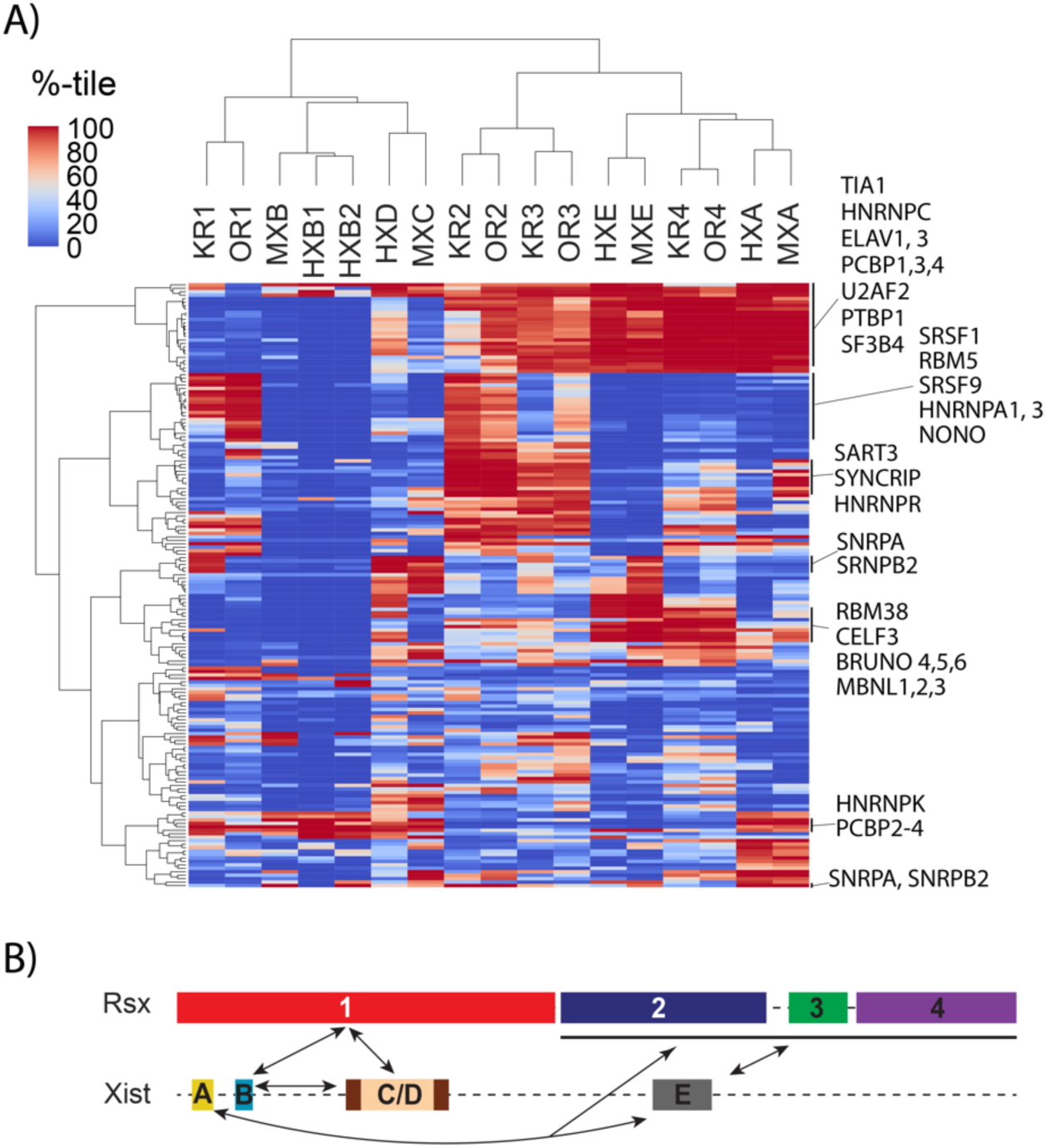
Protein-binding motif enrichment in repeats of *Xist* and *Rsx*, and similarity model. **(A)** Hierarchically clustered heatmap of PWM weighted z-scores for each repeat in *Xist* and *Rsx*, expressed as a percentile relative to the set of all GENCODE M18 lncRNA annotations. **(B)** Regions predicted to have similar protein-binding functions in *Xist* and *Rsx*. Arrows connect domains in each lncRNA that have similar k-mer and motif contents.

Using ranked enrichment of protein-binding motifs as a metric for hierarchical clustering, we identified the same relationships between *Xist* and *Rsx* repeat domains as we did when we hierarchically clustered domains by their k-mer content alone (dendrogram in Figure 6A compared to dendrograms in Figures 2E and 5H). Via protein-binding motif enrichment, *Xist* Repeats B, C, and D formed a second order cluster that next joined with *Rsx* Repeat 1. This clustering order is the same as that detected using k-mer content alone (Figure 6A vs. 2E and 5H). Likewise, protein-binding motif enrichment grouped Repeats A and E together with *Rsx* Repeat 4, while *Rsx* Repeats 2 and 3 formed a separate cluster that joined with the Repeat-A-E-4 cluster. Again, similar clustering patterns were obtained based purely on k-mer content (Figure 6A vs. 2E and 5H). We note that the motifs used to create these clusters are limited in complexity and are capable of recruiting different proteins depending on cellular and sequence contexts (Dominguez et al., 2018; Ray et al., 2013). Thus, the enrichment of a particular protein-binding motif in an individual *Xist* or *Rsx* repeat domain does not provide direct evidence that the protein binds to the domain. Nevertheless, these results are consistent with the notions that lncRNA k-mer content encodes information about protein-binding potential (Kirk et al., 2018), and that the various repeats in *Xist* and *Rsx* encode function through the concerted recruitment of multiple RNA-binding proteins.

A closer inspection of the protein-binding motifs that were enriched in each repeat domain yielded several insights. First, our motif analysis uncovered relationships between repeat domains that were not obvious from direct k-mer comparisons. For example, both human and mouse *Xist* Repeat B were enriched in motifs that recruit polyC-binding proteins and little else (Figure 6A; Table S4). *Rsx* Repeat 1, which most closely resembles *Xist* Repeat B at the level of k-mers, was also enriched in polyC-binding motifs, in both koala and opossum (Figure 6A; Table S4). However, Repeat 1 from koala and opossum *Rsx* were also enriched in many motifs that were absent in Repeat B, such as motifs that bind the proteins SRSF1, SRSF9, and RBM5 (Figure 6A; Table S4). In addition, while both pure k-mer analysis and motif analysis identified similarities between *Xist* Repeats A and E and *Rsx* Repeats 2, 3, and 4, our motif analysis also identified similarities exclusive to pairs of domains within this group, such as similarities between *Rsx* Repeats 2 and 3 and similarities between *Xist* Repeat E and *Rsx* Repeat 4 (Figure 6A; Table S4).

Second, within individual repeat domains, many protein-binding motifs were enriched to extreme levels. Well over half of the motifs analyzed (101 out of 175) were in the 99^th^ percentile in terms of their enrichment relative to other mouse lncRNAs in at least one *Xist* or *Rsx* repeat domain, and all repeat domains in *Xist* and *Rsx* harbored multiple protein-binding motifs that were enriched at the 99^th^ percentile or greater (Figure 6A; Table S4). This extremity was notable considering that most *Xist* and *Rsx* repeats are greater in length than the average mouse lncRNA (Figure 2A). For example, at 13kb in length, *Rsx* Repeat 1 is longer than 99.97% of spliced mouse lncRNAs (Figure 2A). Nevertheless, on a length-normalized basis, multiple protein-binding motifs were enriched in Repeat 1 at the 99^th^ percentile, in both koala and opossum *Rsx*. Inasmuch as motif density is known to be an important driver of associations between proteins and RNA (Dominguez et al., 2018; Kirk et al., 2018; Van Nostrand et al., 2016), our data suggest that at the level of sequence composition, the repeat domains in *Xist* and *Rsx* each have the potential to serve as high-affinity binding platforms for multiple proteins.

Lastly, many of the most strongly enriched motifs in both *Xist* and *Rsx* are known to recruit near-ubiquitous RNA-binding proteins that play core roles in the process of splicing (Wahl and Luhrmann, 2015). These included PTBP1, RMB5, SF3B4, SNRPA, SNRPB2, U2AF2, multiple SR proteins, and multiple HNRNP proteins, including HNRNPA1, HNRNPC, and HNRNPK (Figure 6A; Table S4). We recognize that the motifs available for this analysis are biased towards RNA-binding proteins whose functions are best understood; overwhelmingly, these proteins are splicing factors (Dominguez et al., 2018; Ray et al., 2013). Nevertheless, it is possible that an extreme enrichment for a motif that recruits a ubiquitously expressed splicing factor may confer a function that a single binding motif would not. For example, we presume that the function of Repeat B could not be recapitulated by a single motif that binds HNRNPK (Pintacuda et al., 2017).

## Discussion

*Xist* has served as a paradigmatic regulatory lncRNA for more than 25 years (Balaton et al., 2018; Brockdorff, 2018; da Rocha and Heard, 2017; Sahakyan et al., 2018). Nevertheless, it has been challenging to apply the information gained from the study of *Xist* to other lncRNAs. This is because *Xist* has little linear sequence similarity to other RNAs, even to lncRNAs like *Rsx*, which seem likely to encode analogous functions (Grant et al., 2012; Wang et al., 2014). In the present study, we used a non-linear method of sequence comparison called SEEKR (Kirk et al., 2018) to compare the repetitive regions of *Xist* and *Rsx*. Our data provide sequence-based evidence to support the hypothesis that *Xist* and *Rsx* are functional analogues that arose through convergent evolution, and provide insights into mechanisms through which their repeat domains may encode function.

Unexpectedly, at the level of k-mers, the repeat domains of *Xist* and *Rsx* partitioned into two major clusters. *Xist* Repeats B, C, and D were highly similar to each other and to *Rsx* Repeat 1, whereas *Xist* Repeats A and E were most similar to each other and to *Rsx* Repeats 2, 3, and 4. From prior analyses of sequence content, there is little that would have suggested that the repeats in these two lncRNAs would cluster together in such a manner. However, prior molecular analyses of *Xist* are consistent with such a clustering (Balaton et al., 2018; Brockdorff, 2018; da Rocha and Heard, 2017; Sahakyan et al., 2018).

Specifically, Xist Repeats B and C are known to play important roles in recruiting PRC1 to the inactive X through their ability to bind HNRNPK and possibly other proteins (Pintacuda et al., 2017). The similarity between Repeats B and C and *Xist* Repeat D and *Rsx* Repeat 1 suggests that the latter two repeats may also play roles in recruiting PRC1. Consistent with this possibility, we found that an antibody specific to HNRNPK robustly retrieved *Rsx* RNA in an IP. Moreover, eCLIP data show that *Xist* Repeat D is enriched for HNRNPK binding in human cells (Figure S5; (Van Nostrand et al., 2016)). Thus, even within *Xist*, murid and non-murid mammals may have convergently evolved separate repeats to recruit PRC1, in the form of Repeats C and D, respectively.

Relatedly, *Xist* Repeats A and E have been implicated in recruitment of PRC2 to the inactive X, both via direct and cooperative means (Almeida et al., 2017; Cifuentes-Rojas et al., 2014; Davidovich et al., 2015; Kohlmaier et al., 2004; Ridings-Figueroa et al., 2017; Sunwoo et al., 2017; Wang et al., 2017; Zhao et al., 2008). The similarity between *Xist* Repeats A and E and *Rsx* Repeats 2, 3, and 4 suggests that the *Rsx* repeats could also play roles in recruiting PRC2.

Based on these data, we propose that the two major clusters of repeats in *Xist* and *Rsx* function in part to cooperatively recruit PRC1 and PRC2 to chromatin. Within *Xist*, Repeat B plays a dominant role in recruiting PRC1 via its ability to bind HNRNPK; in turn, PRC1-induced chromatin modifications likely stimulate loading of PRC2 onto chromatin of the inactive X (Almeida et al., 2017; Pintacuda et al., 2017). Nevertheless, a PRC1-dominant model does not preclude other repeats in *Xist* or *Rsx* from functioning in PRC2 recruitment. Indeed, while there does not appear to be a single domain in *Xist* that is absolutely required to recruit PRC2 during the early stages of XCI (Kohlmaier et al., 2004; Wutz et al., 2002), it is possible that multiple domains in *Xist* recruit PRC2 duplicatively, such that deletion of any single domain alone does not cause complete loss in PRC2 recruitment. This hypothesis is supported by prior studies that link both Repeat A and E to recruitment of PRC2 (Almeida et al., 2017; Cifuentes-Rojas et al., 2014; Davidovich et al., 2015; Kohlmaier et al., 2004; Ridings-Figueroa et al., 2017; Sunwoo et al., 2017; Wang et al., 2017; Zhao et al., 2008), and by our own data that show *Xist* Repeats A and E and *Rsx* Repeats 2, 3, and 4 have similar k-mer profiles and motif contents. PRC1, PRC2, and related complexes function cooperatively in flies, mammals, and plants (Blackledge et al., 2015; Li et al., 2018; Schuettengruber et al., 2014). Considering this cooperativity, it is conceivable that the repeat domains in *Xist* and *Rsx* also cooperate to distribute PRC1 and PRC2 on chromatin.

Beyond recruiting PRCs, *Xist* evades nuclear export, it associates with transcribed regions of chromatin, and it induces Polycomb-independent gene silencing (Balaton et al., 2018; Brockdorff, 2018; da Rocha and Heard, 2017; Sahakyan et al., 2018). It is possible that *Rsx* carries out many, if not all of these actions, and that *Rsx* relies on sets of proteins similar to those employed by *Xist* to achieve them (Grant et al., 2012; Wang et al., 2014). We found that all *Xist* and *Rsx* repeat domains harbored extreme levels of enrichment for multiple motifs known to recruit different subsets of RNA-binding proteins. Most of these proteins have been best-characterized in the context of splicing, rather than epigenetic silencing.

In light of these data, we suggest that the repeat domains in *Xist* and *Rsx* may encode some of their functions not by recruiting a set of dedicated RNA silencing factors, but by engaging with ubiquitously-expressed RNA-binding proteins in ways that are distinct from most other RNAs. Such a model was recently proposed (Brockdorff, 2018), and agrees well with what is known about the specificity of RNA-protein interactions. Most RNA-binding proteins have limited sequence specificity, and are capable of binding many thousands of regions in hundreds to thousands of expressed RNAs (Dominguez et al., 2018; Ray et al., 2013; Van Nostrand et al., 2016). SPEN and HNRNPK are two RNA-binding proteins that are critical for *Xist*-induced silencing, yet they clearly associate with RNAs other than *Xist* (Cirillo et al., 2016; Van Nostrand et al., 2016). Relatedly, many other proteins important for *Xist*-induced silencing play central roles in RNA splicing and nuclear export and, through these latter roles, likely associate with a large portion of the transcriptome (Moindrot et al., 2015). Thus, *Xist* and *Rsx* may distinguish themselves from other chromatin-associated transcripts not necessarily by the proteins to which they bind, but by the manner in which they bind these proteins.

That the related repeat domains were present in a different order in *Xist* and *Rsx* supports the notion that within a lncRNA, the order of functional domains is likely to be less important than the presence of the functional domains (Figure 6B). This notion is consistent with a body of work that suggests lncRNAs encode regulatory function in a modular fashion, via discrete domains that recruit distinct subsets of effector proteins (Hacisuleyman et al., 2016; Hezroni et al., 2015; Johnson and Guigo, 2014; Kelley et al., 2014; Kirk et al., 2018; Liu et al., 2017; Lu et al., 2016; Lubelsky and Ulitsky, 2018; Patil et al., 2016; Pintacuda et al., 2017; Smola et al., 2016; Somarowthu et al., 2015; Tsai et al., 2010; Wutz et al., 2002).

From a methodological standpoint, our manuscript outlines approaches that should prove useful in the study of functional domains in other sets of RNAs. Intuitively, k-mer based comparisons like SEEKR seem most likely to succeed in identifying similarity when the domains of interest are repetitive. By nature, repetitive domains that share enrichments of similar subsets of k-mers will be more similar to each other than they will be similar to the average non-repetitive region in the transcriptome.

Nevertheless, similarity between two repetitive domains, when observed, should be carefully considered, especially when the similarity occurs in lncRNAs such as *Xist* and *Rsx*, which are expressed at similar levels in equivalent subcellular compartments (Grant et al., 2012; Wang et al., 2014). Motif density is known to be a dominant factor driving protein/RNA interactions (Dominguez et al., 2018; Kirk et al., 2018; Van Nostrand et al., 2016). All other variables being equal, two lncRNAs that harbor domain-specific k-mer similarity should possess similar protein-binding profiles that could specify similar or analogous function.

However, SEEKR is not limited to analysis of repetitive domains. It also has the ability to detect similarity between repetitive and non-repetitive domains and between strictly non-repetitive domains as well. In any given sequence, a set of k-mers can be arranged in repetitive or non-repetitive ways, and SEEKR has no inherent preference for one over the other. As a contrived example, the sequence of *Xist* Repeat D can be shuffled in a way that eliminates its repeated monomers, yet entirely preserves its k-mer content (File S1). By BLAST, this shuffled sequence has little internal similarity to itself or to Repeat D (Figure S6). Yet, by k-mer content, the shuffled sequence and Repeat D are literally identical (Figure S6). In a real-world example, the top five lncRNAs that SEEKR found to be the most similar to Repeat D are not nearly as repetitive as Repeat D itself (Figure S6). Of all *Xist* and *Rsx* repeats, Repeat D is the most complex (Figure 2B). Nevertheless, these results demonstrate that k-mer based similarity searches performed with repetitive domains can identify non-repetitive top hits.

With regard to non-repetitive domains, our 2018 study showed that SEEKR rivaled BLAST-like alignment in its ability to detect lncRNA homologues in human and mouse (Kirk et al., 2018). The majority of homologues detected by SEEKR either lacked obvious repetitive elements, or were predominantly comprised of non-repetitive sequence; the lncRNAs *H19, Hottip*, *Malat1*, *Miat*, and *RMST* being specific examples. We also found that SEEKR could identify *Xist*-like repressive activity in several synthetic and natural lncRNAs that lacked repetitive elements (Kirk et al., 2018). Thus, even in non-repetitive regions of RNA, SEEKR should be capable of detecting meaningful similarities. However, functional domains comprised of high-complexity sequence elements will likely remain challenging to identify, regardless of the method in use.

Key variables to decide upon when using SEEKR are the k-mer length and the appropriate set of RNAs that define the background k-mer frequency; i.e. the set of RNAs used to define the means and standard deviations from which k-mer z-scores are calculated. At present, data regarding functional domains in lncRNAs are too limited to arrive at conclusive recommendations for either variable. We favor using a k-mer length at which 4^k most closely resembles the length of the shortest domain being analyzed. This approach minimizes the number of k-mers that yield counts of zero in the domain. Data from the present study as well as our prior work suggest that this minimization increases discriminatory power (Figure S4; (Kirk et al., 2018)).

In terms of the set of RNAs that should be used to define the background k-mer frequency, it is worth noting that SEEKR measures relative, not absolute, similarity. Pearson’s r values returned by SEEKR reflect the similarity between two sequences relative to the k-mer frequency present in the background set of RNAs. We have found that using a background set of all lncRNAs in a genome provides a convenient way to identify trends. For example, in the present study, we used all known spliced lncRNAs in the mouse as a background set. Accordingly, we were able to identify properties in the repeat domains of *Xist* and *Rsx* that were distinct from the average spliced lncRNA annotated by GENCODE (Derrien et al., 2012).

In our initial description of SEEKR, we used k-mer contents of full-length lncRNAs as search features; we did not examine k-mer contents at the level of individual domains (Kirk et al., 2018). The domain-centric approaches outlined in the present study may be better suited for lncRNAs such as *Xist* and *Rsx*, which have multiple functions that are likely to be distributed amongst multiple domains. Indeed, at the level of k-mers, full-length *Xist* and *Rsx* were negatively correlated to each other. Similarities between the two lncRNAs emerged only when we took a domain-centric approach. Other eutherian lncRNAs known to harbor *Xist*-like silencing function, such as *Kcnq1ot1* and *Airn*, are exceptionally long – each on the order of 90kb. Extrapolating from our findings above, we would expect these lncRNAs to harbor the greatest levels of similarity to each other not at the level of their full-length transcripts, but at the level of specific domains.

## Acknowledgements

We thank UNC colleagues for discussions. This work was supported by National Institutes of Health (NIH) Grant GM121806 and Basil O’Connor Award #5100683 from the March of Dimes Foundation (to J.M.C.), NIH Grant R014214 (to P.B.S.), and the Australian Research Council grant DP180100931 (to P.D.W.). D.S. was supported in part by an NIH training grant in pharmacology (T32 GM007040). J.R.W. was supported by the University Cancer Research Fund.

## Methods

### Dot plots

Dot plots were generated using EMBOSS dotmatcher (Rice et al., 2000). For clarity, different visualization thresholds were used to generate the different dot plots shown in the manuscript. Figure 1A, C, and D, and Figure 5A and B, used a window size 10 and a threshold of 40. Figure 1B, E, and F used window size 10 and threshold 45. Figure S1 used a window size 20 and a threshold of 50.

### Definition of repeat domains in *Xist* and *Rsx*, and non-human/non-mouse *Xist* sequences

The sequence of all *Xist* and *Rsx* repeat domains used in this work can be found in File S1. The sequences of all full-length *Xist* and *Rsx* lncRNAs used in this work can be found in File S2. The spliced mouse *Xist* sequence was sourced from the mm10 build of the mouse genome and annotations for the tandem repeats were sourced from (Brockdorff et al., 1992). The spliced human *Xist* sequence was sourced from the hg38 build of the human genome and the annotations for the tandem repeats were sourced from (Brown et al., 1992; Yen et al., 2007).

The sequences of spliced *Xist* used to generate the dot plots in Figure S1 were obtained directly from annotations in the UCSC genome browser (Tyner et al., 2017), or, for genomes in which full annotations were unavailable, were reconstructed from partial annotations by UCSC and RNA-seq data from (Hezroni et al., 2015). In the case of the vole *Microtus rossiaemeridionalis*, *Xist* sequence was obtained directly from (Nesterova et al., 2001).

Spliced koala *Rsx* was obtained from (Johnson et al., 2018). To identify repeat domains, *Rsx* was aligned to itself using EMBOSS dotmatcher with a 10bp window and a 40% threshold (Rice et al., 2000). Starts and stop positions of each repeat were defined by visual inspection of the dot plot. We considered separating the fourth major repeat in *Rsx* into two repeat domains, one 500bp and the other 5000bp in length (see Figure 1D); however, analysis of the shorter sub-repeat within Repeat 4 revealed its k-mer content to be highly similar to the larger sub-repeat (not shown). Thus, to simplify our analyses and to clarify our presentation, we elected to merge the sub-repeats. Repeat domains in opossum *Rsx* (after filling in the gaps in assembly; see below) were defined in the identical manner.

### K-mer based comparisons (i.e. SEEKR)

SEEKR was performed essentially as described in (Kirk et al., 2018), with minor modifications. As a reference for normalization, we first calculated the mean and standard deviation for all k-mers at k = 4 in the GENCODE M18 lncRNA annotation file. We then generated length normalized counts of all k-mers at k = 4 for each repeat domain in *Xist* and *Rsx* and calculated z-scores for each k-mer by subtracting the mean and dividing by the standard deviation for each k-mer from our reference set of GENCODE lncRNAs. Prior to performing Pearson’s correlation, z-scores were log_2_ transformed.

To generate the distributions of Pearson’s values in Figure 2B and Figure 5I, we calculated the k-mer profile for each repeat domain and each GENCODE M18 lncRNA using the mean and standard deviation values from the full-length GENCODE M18 lncRNA annotation file, as described above. We then log_2_-transformed the z-scores and used Pearson’s correlation to compare all lncRNAs to the *Xist* repeat in question.

### Hierarchical clustering

Hierarchical clustering was performed using the scipy hierarchy package in Python 3.6 (Jones et al., 2001), with distance defined as *d* = 1 − *r*, where *r* is defined as the Pearson correlation, using complete linkage.

### *De novo* motif analysis

Motifs in each *Xist* and *Rsx* repeat domain were detected with MEME (version 5.0.2; (Bailey et al., 2009)), run using the following options: -mod anr -dna -bfile bkg.meme -nmotifs 100 -minw 4-maxw 12 -maxsites 1000, where the “bkg.meme” file specified a background frequency of 0.25 for all four nucleotides.

### Consecutive k-mer analyses

To calculate the sums of z-scores for k-mers containing matches to mononucleotide runs in Figures 4A-D, we used the following approach. A mononucleotide run was defined as at least two consecutive occurrences of the nucleotide in question. For each nucleotide [A|C|G|T], we multiplied the z-score for each k-mer that contained a run by (the nucleotide length of the run minus 1). The sum of these products for each repeat domain at k-mer length k = 5 is plotted in Figures 4A-D. Identical trends were seen using k-mer lengths k = 4, 5, and 6 (Figure S3). K-mer length k = 5 was chosen for plotting in Figure 4 to emphasize trends that were present but less pronounced when using k-mer length k = 4. The set of mouse lncRNAs from GENCODE M18 was used to derive z-scores that described the length normalized abundance of each k-mer in each repeat domain.

### Weighting k-mer z-scores by likelihood of matching the HNRNPK-binding motif

To weight the sums of z-scores by the HNRNPK PWM in Figure 4E we performed the following calculation. For all k-mers at k = 5 we calculated the probability of a given k-mer’s sequence occurring in the PWM for HNRNPK. The probability was defined as the independent probability of each letter in the k-mer occurring at the corresponding location within the PWM for each possible frame within the PWM. The HNRNPK motif is 8nt long, therefore there were 3 possible frames for a 5-mer to fall within. The z-score for the k-mer in question was then weighted by taking the sum of the product between the z-score and each probability. The height of the bars in Figure 4E represent the sum of weighted z-scores for each *Xist* and *Rsx* repeat domain. The set of mouse lncRNAs from GENCODE M18 was used to derive z-scores that described the length normalized abundance of each k-mer in each repeat domain.

### Detecting HNRNPK-binding motif matches

Motifs occurrences in each *Xist* and *Rsx* repeat domain were detected with FIMO (version 5.0.2; (Bailey et al., 2009)), run using the following non-default option: --thresh 0.01.

### RNA Immunoprecipitation

Cultured female *M. domestica* fibroblast cells were harvested at 70% confluency by scraping, then aliquoted into 1 × 10^7^ cells, pelleted by centrifugation at 200g, then snap-frozen and stored at −80°C until used. RIPs from non-crosslinked cells were performed essentially as described in (Zhao et al., 2010), using the following antibodies from Abcam: H3K27me3 (ab6002), CTCF (ab70303), HNRNPK (ab39975), and mouse IgG (ab18413). Briefly, cell pellets were gently resuspended in 1 mL of ice-cold RIPA buffer supplemented with 1× EDTA-free Proteinase Inhibitor Cocktail (Thermo Scientific) and lysed for 15 min at 4°C. Samples were sonicated at 4°C (Qsonica Q700 with cup horn accessory) at 12% amplitude for fifteen 30 second intervals, with 30 second resting steps between intervals. Cell debris was removed by centrifugation (at 6000 g for 5 minutes), and samples were subsequently diluted to 1mg of protein per ml with ice-cold RIPA buffer. Lysates with 1mg of total protein (i.e. 500ul) were incubated with the appropriate antibody coupled to Protein G beads (Life Technologies), overnight at 4 °C with end-over-end rotation. Beads with no antibodies (mock IP) were used as background control. Beads were removed from lysate using a magnetic stand and were re-suspended in 1ml of ice cold NP-40 buffer (50 mM Tris at pH 7.5, 50 mM NaCl, 10 mM EDTA, 1% Nonidet P-40, 0.5% sodium deoxycholate, 0.1% SDS) and washed for 15min at 4 °C with end-over-end rotation, repeated twice, followed by three washes with RIPA buffer. Following the last wash, beads were collected and re-suspended in 1ml of Trizol (Life Technologies) for RNA extraction. 10% of the input lysate (i.e. 50ul) was processed in parallel. RNA was cleaned using RNeasy spin columns (Qiagen), following the manufacturer’s “RNA Cleanup” protocol, with on-column RNase-free DNase Set (Qiagen) treatment. cDNA was synthesized using input and immunoprecipitated RNA with SuperScript III reverse transcriptase (Life Technologies) and random hexamer priming. *Rsx* was detected by RT-qPCR (in technical triplicate) with primer pair L2 from (Grant et al., 2012). Cycle threshold (Ct) values were normalized to input and relative to the IgG. Fold enrichment was determined by relative quantification, which was calculated using the 2^e(−ΔΔ Ct) method. The level of Gapdh mRNA enrichment was used as an internal non-target index in the qPCR analysis.

### Nanopore Sequencing and annotation of opossum *Rsx*

High molecular weight DNA from VMRC18-839J22 and VMRC18-303M7 BACs was prepared using the NucleoBond BAC 100 kit (Machery Nagel). DNA from the two BAC preparations was pooled, sheared to an average length of 20kb using a g-TUBE (Covaris), and then sequenced on the Oxford Nanopore Technologies (ONT) MinION using an R9.4 flow cell (FLO-MIN106) following the 1D ligation protocol (SQK-LSK109).

Reads were base-called with Albacore 2.3.1 (ONT) then assembled using Flye 2.3.5b (Kolmogorov et al., 2018). The six resulting scaffolds were aligned to *E. coli* K12 (NC_000913.3), opossum chromosome X (MonDom5, NC_008809.1) and the pCC1BAC cloning vector (EU140750.1). Scaffolds consisting entirely of *E. coli* or cloning vector DNA were removed. Three scaffolds aligned to adjacent regions of the MonDom5 X chromosome. These were merged together into a single candidate assembly sequence that was then polished iteratively with Racon 1.3.2 four times (Vaser et al., 2017), followed by Nanopolish 0.10.1 (Loman et al., 2015), to produce a final complete assembly of 235,139 nucleotides (File S3).

This polished assembly sequence was aligned again to MonDom5 using BLASTN to establish start and end coordinates to use as a reference when replacing the gaps in MonDom5 with the completed sequence in our assembly. The final sequence of opossum *Rsx* used in this work (File S2) was generated using splice annotations from (Grant et al., 2012), and replacing the N’s in mondom5 with the corresponding sequence from our polished assembly (nucleotide substitutions are listed in Table S3). Raw sequencing reads were deposited in NCBI’s SRA, under accession number PRJNA522427.

### Shuffling of Repeat D sequence

The sequence of Repeat D was shuffled using ushuffle (Jiang et al., 2008).

---

**Figure S1.**
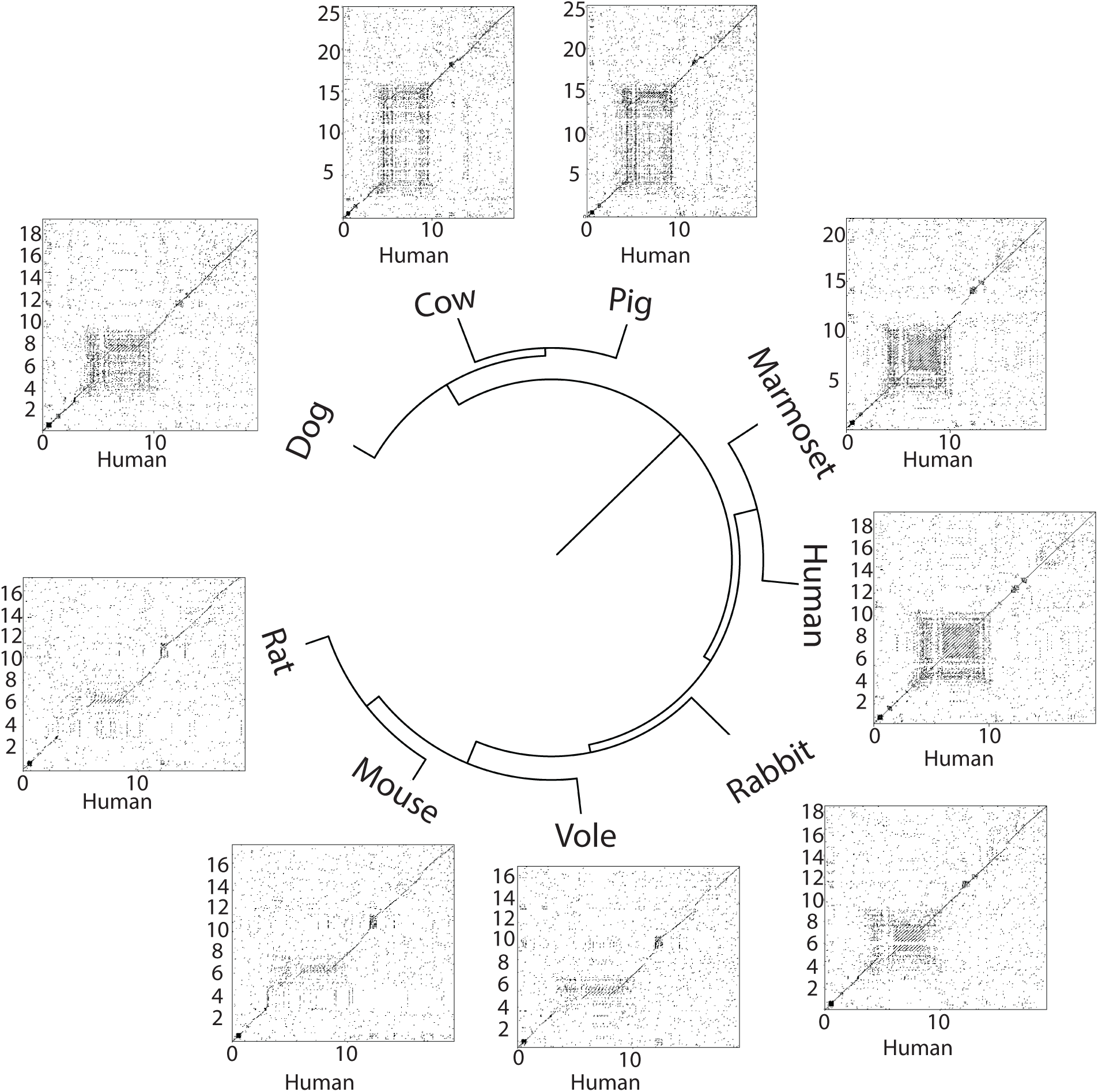
Conservation of Repeat D-like sequence in non-murid mammals. Human *Xist* aligned against *Xist* from other mammals using dotmatcher with window size = 20 and threshold = 50. Human *Xist* is along the x-axis and the indicated species is along the y-axis. Repeat D-like regions tend to be the largest domains of similarity between human and non-murid *Xists*.

**Figure S2.**
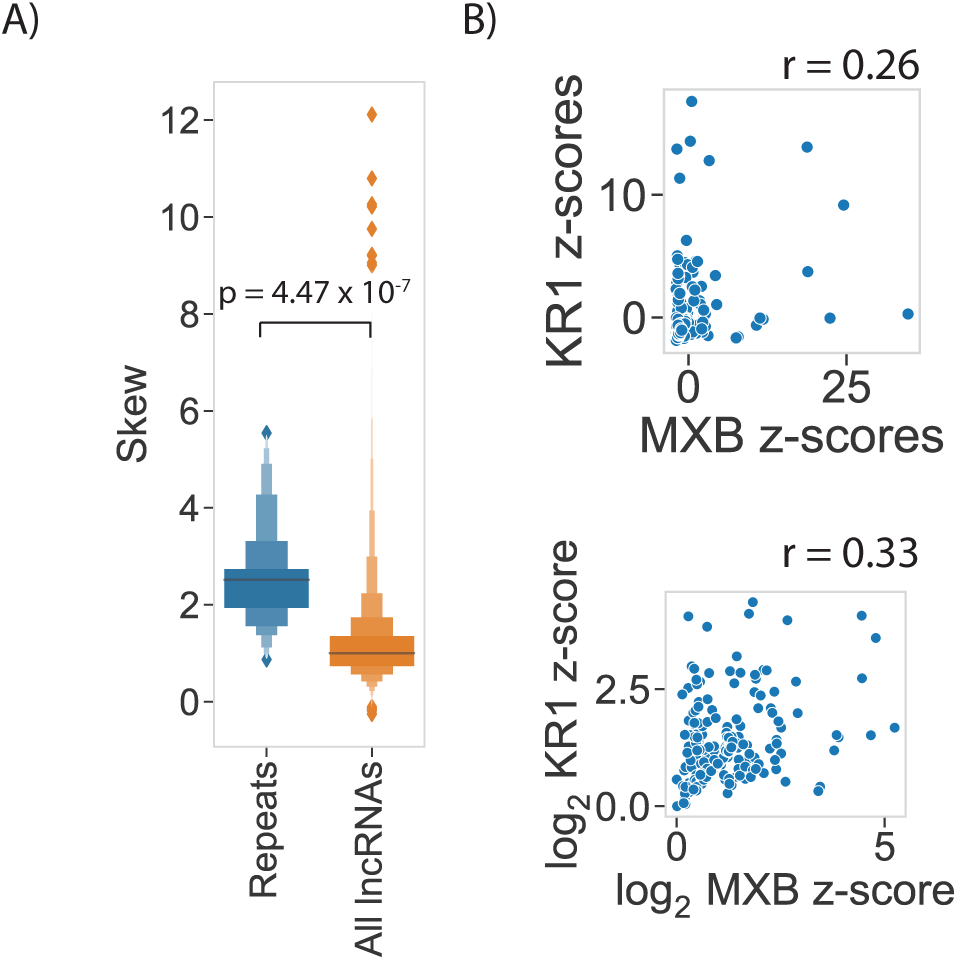
Log-log transformation of z-scores improves correlation between tandem repeat domains. **(A)** *Xist* and *Rsx* repeats (“Repeats”) and the set of reference GENCODE lncRNAs have z-score vectors with positive (right) skew; z-scores in *Xist* and *Rsx* repeats are significantly more skewed than those in the reference set of GENCODE lncRNAs (Mann-Whitney U Test). **(B)** Representative plots of improvement in correlation from untransformed (top) and log_2_-log_2_ transformed z-scores (bottom). Repeat domain abbreviations are the same as those in Figure 2.

**Figure S3.**
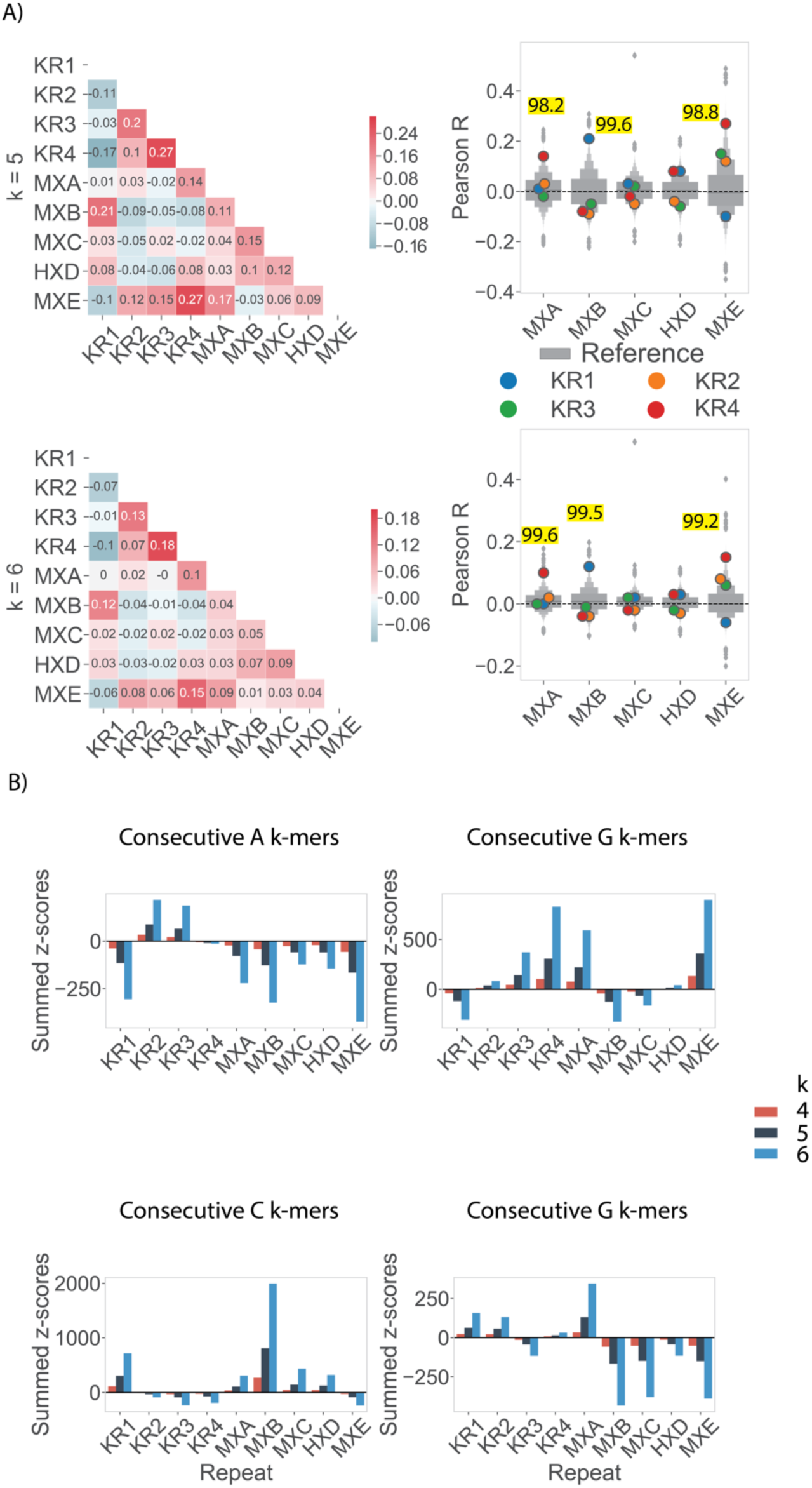
Similarities between *Xist* and *Rsx* are apparent at multiple k-mer lengths. **(A)** Similarities between repeat domains in *Xist* and *Rsx*, as plotted in Figure 2, for k-mer lengths k = 5 and 6. **(B)** Enrichment in mononucleotide runs in the repeat domains of *Xist* and *Rsx*, as plotted in Figure 4, for k-mer lengths k = 4, 5, 6.

**Figure S4.**
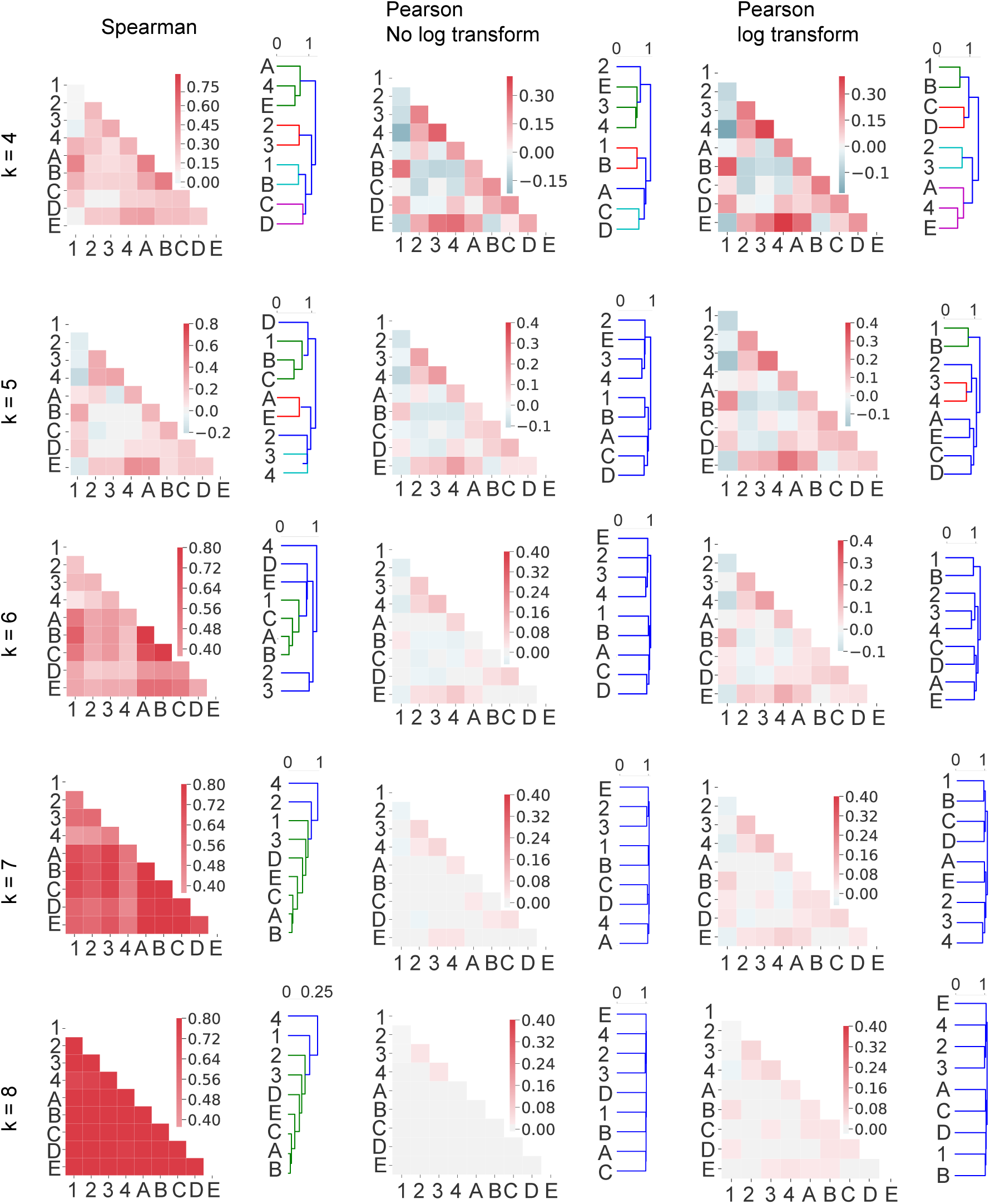
Different correlation metrics identify similar relationships between *Xist* and *Rsx* repeat domains. A direct comparison of *Xist* and *Rsx* domain similarities detected using Spearman’s correlation of untransformed k-mer counts, Pearson’s correlation of untransformed k-mer counts, and Pearson’s correlation of log2-transformed untransformed k-mer counts. Dendrogram colors represent clusters for all descendent links beneath the first node in the dendrogram with distance less than 70% of the maximum distance between all clusters.

**Figure S5.**
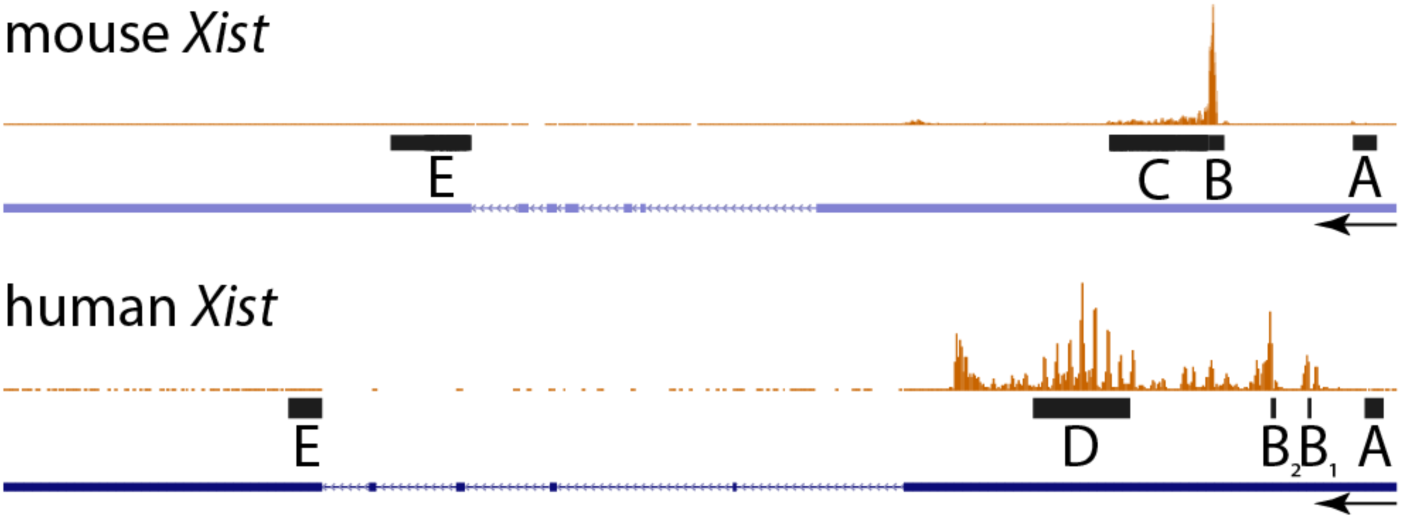
*Xist* Repeats B, C, and D are enriched for HNRNPK-binding. UCSC wiggle-density display of HNRNPK CLIP data (orange) aligned over the mouse (mm9) and human (hg38) *Xist* genomic loci. Mouse and human clip data are from (Cirillo et al., 2016; Van Nostrand et al., 2016). Arrows denote direction of *Xist* transcription. Black rectangles indicate genomic locations of mouse and human repeat sequences used in this work.

**Figure S6.**
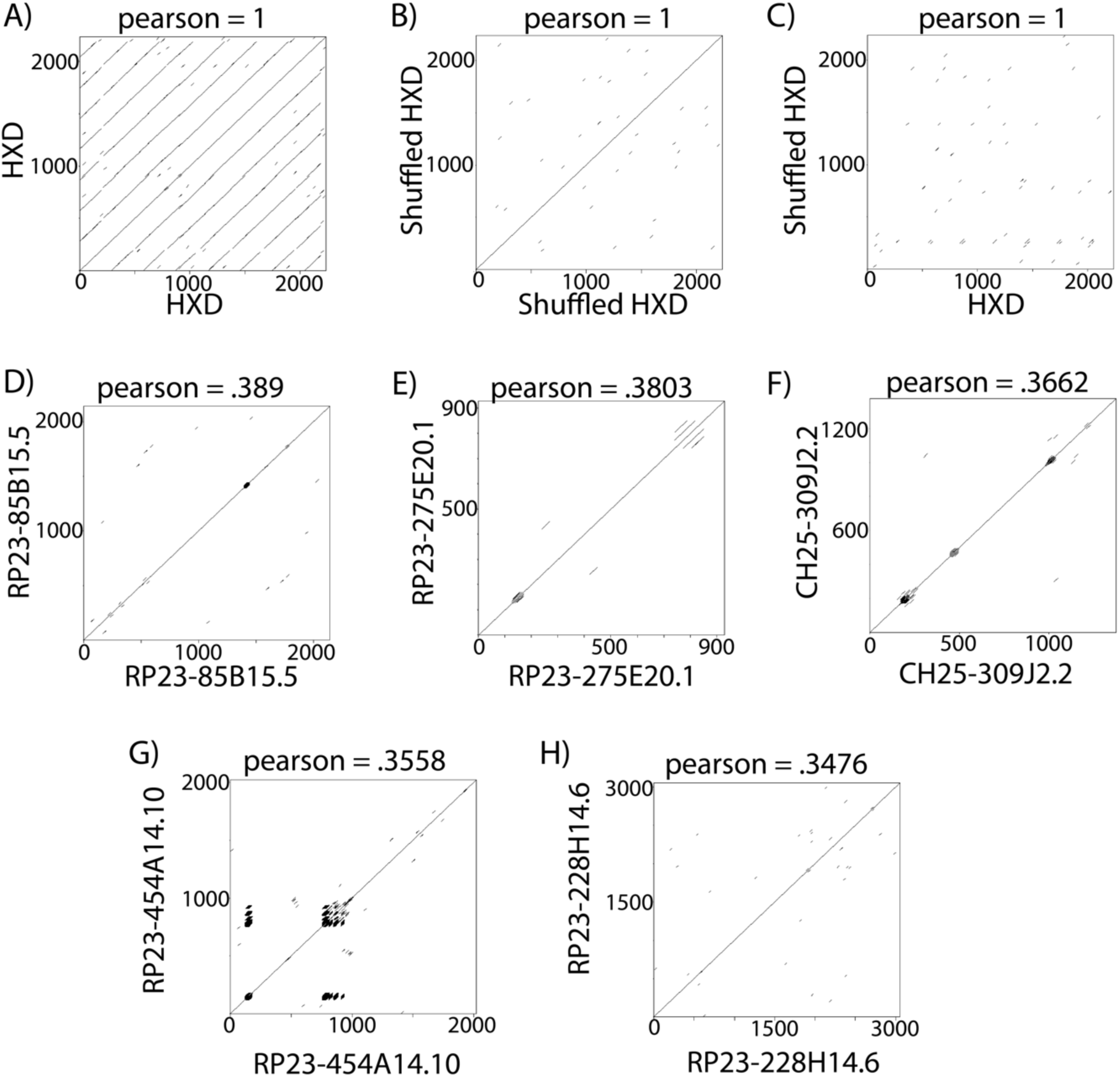
Dot plot alignments and SEEKR-defined similarity between human *Xist* Repeat D, shuffled Repeat D, and the top 5 mouse lncRNAs that are most similar to Repeat D. Repeat D was shuffled using uShuffle and preserving k-mer content at k = 4 (Jiang et al., 2008). Dotplots were generated using a window size 20 nucleotides and a threshold of 50% identity.

**File S1. A file containing the sequence of all repeat domains used in this work, in fasta format.**

**File S2. A file containing the sequence of full-length *Xist* and full-length *Rsx* lncRNAs used in this work, in fasta format.**

**File S3. A file containing the ungapped sequence assembled from Nanopore sequencing of DNA from *M. domestica* BACs VMRC18-839J22 and VMRC18-303M7, in fasta format.**

**Table S1. Percentiles of *Rsx* Repeat Domains.** The percentile of similarity between *Rsx* repeats and *Xist* repeats relative to all GENCODE M18 spliced lncRNAs, as reported in Figure 2D and in Figure 5I.

**Table S2. Top 1000 most similar lncRNAs to the different repeat domains in *Xist*.** Ranked lists of the top 1000 most similar lncRNAs to the different repeat domains in *Xist*. *Xist* and koala *Rsx* repeat domains are also added to each list.

**Table S3. Location of Sequencing Gaps in mondom5 *Rsx.*** 5′ and 3′ alignments of the BAC to the mondom5 genome build are annotated first. Coordinates of ‘N’ nucleotides in the genome build (left) and coordinates within our BAC of sequences aligning to the left and right of each ‘N’ in mondom5, the length of sequence in replacing the ‘N’ in mondom5, and the difference in sequence length are annotated next (right).

**Table S4. Percentile enrichment of protein-binding motifs in the same order as presented in** Figure 6A. PWM weighted z-score sums for each *Xist* and *Rsx* repeat converted to a percentile relative to the GENCODE V18 mouse lncRNA annotation file. The rows, corresponding to each PWM, and columns, corresponding to *Xist* and *Rsx* repeats, are sorted using hierarchical clustering, with euclidean distance and Ward linkage. Proteins corresponding to each PWM are listed in the final column.

